# Elucidating the role of muscarinic acetylcholine receptor (mAChR) signaling in efferent mediated responses of vestibular afferents in mammals

**DOI:** 10.1101/2023.07.31.549902

**Authors:** Anjali K. Sinha, Choongheon Lee, Joseph C. Holt

## Abstract

The peripheral vestibular system detects head position and movement through activation of vestibular hair cells (HCs) in vestibular end organs. HCs transmit this information to the CNS by way of primary vestibular afferent neurons. The CNS, in turn, modulates HCs and afferents via the efferent vestibular system (EVS) through activation of cholinergic signaling mechanisms. In mice, we previously demonstrated that activation of muscarinic acetylcholine receptors (mAChRs), during EVS stimulation, gives rise to a slow excitation that takes seconds to peak and tens of seconds to decay back to baseline. This slow excitation is mimicked by muscarine and ablated by the non-selective mAChR blockers scopolamine, atropine, and glycopyrrolate. While five distinct mAChRs (M1-M5) exist, the subtype(s) driving EVS-mediated slow excitation remain unidentified and details on how these mAChRs alter vestibular function is not well understood. The objective of this study is to characterize which mAChR subtypes drive the EVS-mediated slow excitation, and how their activation impacts vestibular physiology and behavior. In C57Bl/6J mice, M3mAChR antagonists were more potent at blocking slow excitation than M1mAChR antagonists, while M2/M4 blockers were ineffective. While unchanged in M2/M4mAChR double KO mice, EVS-mediated slow excitation in M3 mAChR-KO animals were reduced or absent in irregular afferents but appeared unchanged in regular afferents. In agreement, vestibular sensory-evoked potentials (VsEP), known to be predominantly generated from irregular afferents, were significantly less enhanced by mAChR activation in M3mAChR-KO mice compared to controls. Finally, M3mAChR-KO mice display distinct behavioral phenotypes in open field activity, and thermal profiles, and balance beam and forced swim test. M3mAChRs mediate efferent-mediated slow excitation in irregular afferents, while M1mAChRs may drive the same process in regular afferents.

## Introduction

The vestibular system is responsible for our general sense of balance, detecting head motion, and providing critical cues for gaze stabilization and spatial navigation (Barany, 1906; Goldberg & Fernandez, 1975; Highstein, 1991; Hitier et al., 2014; Hufner et al., 2007; Jacob et al., 2014; Mao et al., 2021; Nguyen et al., 2021; Wallace et al., 2002). The encoding of head position and movement begins in the inner ear with the activation of hair cells in vestibular end organs. These hair cells transmit this information to the CNS by way of primary vestibular afferent neurons. The CNS can, in turn, modulate this process through the activation of the efferent vestibular system (EVS). In mammals, cholinergic EVS neurons arise bilaterally from e-group nuclei in the dorsal brainstem, and they project both contralaterally and ipsilaterally to synapse onto type II hair cells and primary vestibular afferents (Gacek and Lyon 1974; Warr 1975; (Goldberg & Fernandez, 1980; Jordan et al., 2013; Leijon & Magnusson, 2014; Lorincz et al., 2021; Mathews et al., 2017; Mathews et al., 2015; Perachio & Kevetter, 1989; Purcell & Perachio, 1997).The anatomy and projections of EVS neurons are reasonably well-established in mammals and recent studies have provided further insight into some of the signaling mechanisms utilized by the EVS to modulate hair cell and afferent electrophysiology.

Electrical stimulation of cholinergic EVS neurons in mammals gives rise to at least three types of kinetically- and mechanistically-distinct responses in vestibular afferents including a fast excitation, fast inhibition, and slow excitation (Brichta & Goldberg, 2000; Goldberg & Fernandez, 1980; Jordan et al., 2013; Lee et al., 2021; Marlinski et al., 2004; McCue & Guinan, 1994; Schneider et al., 2021). As the names imply, fast excitation and fast inhibition peak and decay within hundreds of milliseconds of EVS stimulation onset and termination, while slow excitation takes several seconds to peak after stimulation onset and persist for tens of seconds following stimulus termination. It has been proposed that activation of a4b2-containing nicotinic acetylcholine receptors (α4β2*-nAChRs) on vestibular afferents leads to EVS-mediated fast excitation (Holt et al., 2015; Schneider et al., 2021) while the sequential activation of α9α10-nAChRs and calcium-activated, small-conductance potassium (SK) channels in type II hair cells drives the EVS-mediated fast inhibition (Holt et al., 2015; Poppi et al., 2018; Yu et al., 2020). In contrast, slow excitation relies on the activation of muscarinic AChRs (mAChRs) and the subsequent closure of potassium channels in vestibular afferent neurons (Holt et al., 2017; Perez et al., 2009; Poppi et al., 2020; Ramakrishna et al., 2021; Schneider et al., 2021). However, the specific mAChR subtype(s) driving EVS-mediated slow excitation has not been identified.

The mAChR family consist of five subtypes and belong to the superfamily of metabotropic (aka G-protein coupled) seven transmembrane receptors (Caulfield & Birdsall, 1998; Eglen, 2005; Thiele, 2013; Wess, 2004; Wess et al., 2007). The mAChRs are divided into two major subgroups based on their preferential effector G-protein and downstream signaling cascade. Even-numbered mAChRs (M2/M4) use the G-protein Gi/o to inhibit adenylyl cyclase and decrease levels of cAMP which reduces protein kinase A (PKA) activity and cAMP-dependent effectors (Wess, 2004). Odd-numbered mAChRs (M1/M3/M5) primarily utilize the G-protein Gq/11 to activate phospholipase C-β (PLC-β) to hydrolyze phosphatidylinositol 4,5–bisphosphate (PIP2) in the membrane to generate diacyl glycerol (DAG) and inositol 1,4,5-triphosphate (IP3). IP3 released from the membrane binds to IP3-receptors (IP3R) on the ER membrane and releases calcium from internal stores (Felder, 1995). Elevated intracellular calcium and DAG work together to activate protein kinase C (PKC) that can phosphorylate downstream substrates. Additionally, activation of odd-numbered mAChRs can also close a potassium conductance called the M-current. Closure of the M-current will increases input impedance and cellular excitability. Both *in vivo* and *ex vivo* studies have demonstrated a role for M-currents in EVS-mediated slow excitation of vestibular afferents which suggests that odd-numbered mAChRs are involved (Holt et al., 2017; Perez et al., 2009; Perez et al., 2010; Ramakrishna et al., 2021; Schneider et al., 2021). While EVS-mediated slow excitation can be blocked by using nonselective mAChR antagonists like scopolamine, atropine, and glycopyrrolate (Lee et al., 2021; Schneider et al., 2021), the specific mAChR subtype(s) that mediate EVS-mediated slow excitation have not been examined.

Previous work in the field using autoradiograph labelling with the nonselective mAChR antagonist QNB and immunolabelling have suggested the presence of mAChRs at various locations within the vestibular neuroepithelium, including HCs and afferents (Drescher et al., 1999; Li et al., 2007). Additionally, in-situ hybridization and PCR studies indicate the presence of all five mAChR subtypes in the peripheral vestibular system (Ishiyama et al., 1997; Wackym et al., 1996). While electrophysiological and pharmacological evidence make the case for mAChRs on hair cells and afferents (Derbenev et al., 2005; Guo et al., 2012; Holman et al., 2019; Holt et al., 2017; Li & Correia, 2011) these studies did not further characterize any of the underlying mAChR subtypes.

In the current study, we evaluated the role of different mAChR subtypes in EVS-mediated slow excitation of vestibular afferents. In anesthetized mice, we used sharp glass microelectrodes to record action potentials from vestibular afferents leaving the inner ear while stimulating EVS neurons in the brainstem with platinum-iridium electrodes. We then used various mAChR-subtype selective drugs to characterize the mAChRs involved in EVS-mediated slow excitation of vestibular afferents. We found that the M3mAChR selective antagonists were more effective at blocking EVS-mediated excitation compared to either M1- or M5mAChR selective compounds, suggesting M3mAChR as the main regulator of EVS-mediated slow excitation. M3mAChR knockout (KO) mice exhibited absent or attenuated EVS-mediated slow excitation in irregularly-firing vestibular afferents, while EVS-mediated slow excitation in regularly-firing vestibular afferents appeared to be unaffected. The distribution of resting afferent discharge rates versus CV*, a measure of discharge regularity, were similar across M3mAChR controls and knockout animals. Similarly, M3mAChR knockout (KO) mice exhibited normal vestibular sensory-evoked potentials (VsEPs) whose amplitude, threshold, and latency metrics were not significantly different from control animals. In conjunction with normal discharge rates, the VsEP data suggests that irregularly-firing vestibular afferents exhibit normal function despite the loss of EVS-mediated slow excitation. In contrast, systemic administration of the general mAChR agonist, oxotremorine-M, enhanced VsEP amplitudes, decreased VsEP threshold and decreased VsEP latencies in control mice, but not in M3mAChR-KO mice, consistent with the loss of EVS-mediated slow excitation in irregularly-discharging afferents. Finally, M3mAChR-KO mice showed deficits in various vestibular related behaviors including open-field activity, balance beam, swim test, and vestibulo-autonomic thermal responses.

## Materials and Methods

### Animals

All animal procedures were performed in accordance with NIH’s Guide for Care and Use of Laboratory Animals and approved by the University Committee for Animal Resources (UCAR) at the University of Rochester Medical Center (URMC). *<u>Control mice</u>*: C57BL/6J WT mice of both sexes were obtained from Jackson Laboratory and housed in 1-way rooms in the URMC vivarium under standard 12-h light:dark cycle and free access to food and water. Experiments in C57BL/6J mice were performed in animals weighing 20-35 g and 49-180 days. *<u>Transgenic mice</u>*: M3mAChR knockout (KO) mice (Yamada et al., 2001) and M2/M4mAChR double-knockout (DKO) were obtained from Dr. Jurgen Wess at NIDCD and bred in-house within the URMC vivarium. Experiments in mAChR-KO animals were performed in animals weighing 15-30 g and aged 70-150 days. Tail snips were sent to Transnetxy to confirm genotype of the mice.

### Animal preparation

Mice were either anesthetized with IP urethane/xylazine (1.6g/kg; 20mg/kg) or tribromoethanol (TBE, aka Avertin; 250mg/kg). Once anesthetized, a tracheotomy was performed for intubation using PTFE tubing (1.5 cm long, 1mm diam.) subsequently secured with surgical thread ligatures (Moldestad et al., 2009). The free end of the PTFE tubing was connected to a mechanical ventilator set at 100 bpm (model 683, Harvard Apparatus). Body temperature (∼37.5 °C) was maintained using a heating pad and closed-loop homeothermic monitoring system while heart rate was monitored using a 3-lead EKG and differential amplifier (Warner Instrument, Holliston, MA). The mouse’s head was secured using a stereotaxic apparatus. After removal of the dorsal scalp and posterior neck muscles, sections of the rear skull were removed with micro-rongeurs to expose the cerebellum and inferior colliculi. The right cerebellar hemisphere, flocculus, and parafloculus were aspirated to expose the arcuate eminence and the common crus. The cerebellar vermis was aspirated using a 5Fr Frazier suction exposing the floor of the 4th ventricle (Lee et al., 2021; Schneider et al., 2021).

### Afferent recordings

To record extracellular spike activity from spontaneously-discharging vestibular afferents, sharp borosilicate microelectrodes (BF150-86-10, Sutter Instruments) with impedance of 60-120 MOhms were filled with 3M KCl and mounted on a 3-axis micromanipulator to be slowly advanced into the superior division of CNVIII just as it exits the otic capsule. An Ag/AgCl Teflon coated wire was inserted into the microelectrode and connected to a preamplifier headstage (Biomedical Engineering, Thornwood, NY). Prior to recording, residual voltage was zeroed, microelectrode capacitance was neutralized by driving the shield of the input cable, and electrode resistance was balanced using dial settings on the amplifier.

### Efferent stimulation

To stimulate EVS neurons in the brainstem, we used a custom linear 4-lead platinum-iridium array. The EVS electrode was inserted into the floor of the 4th ventricle at the midline with the first electrode lead located just caudal to the facial colliculi (Schneider et al., 2021). Efferent stimuli were delivered from a stimulus isolator (model A360; World Precision Instruments, Sarasota, FL, United States) whose timing was managed using in-house Spike2 scripts on a PC with a micro1401 interface (Cambridge Electronic Design). Standard EVS stimuli consisted of 5-s trains of 100-150µsec constant current shocks delivered at 333 shocks/s (Brichta & Goldberg, 2000; Goldberg & Fernandez, 1980; Lee et al., 2021; Schneider et al., 2021). We adjusted shock pulse amplitude to determine the threshold (T, 20-50μA) and maximum currents (40-250 μA) while avoiding antidromic activation of afferent fiber. Inter-train intervals of 60s or longer were provided to permit afferent discharge rates to return to pre-stimulus values before the next shock train was given.

### Data acquisition

Data acquisition was also managed using in-house Spike 2 scripts on a PC with micro1401 interface (Cambridge Electronic Design). Afferent voltage signals were low-pass filtered (1KHz, four-pole Bessel; wavetrek) and sampled at 10 KHz. Spike2 data files were analyzed using custom macros in IgorPro 6.36 (WaveMetrics). Stimulation artifacts were removed offline by subtracting an averaged shock artifact from each shock stimulus in the raw data.

### Drug Administration

We have previously demonstrated that various cholinergic drugs can be effectively administrated in one of three ways to reach efferent synapses in the inner ear of mice (Lee et al., 2021). These same administration routes were used in this study and include, in order of increasing procedural complexity: (1) Systemic administration using standard intraperitoneal (IP) injection, (2) Delivery into middle ear space via intrabullar administration (IB) or (3) slowly injecting into the perilymph through an intracanal approach (IC). Briefly, for IB injections, an incision was made behind the right pinna and bulla was exposed by removing the muscle above it. A small hole was made in the otic bulla using the tip of 30-G needle to accommodate plastic tubing (100-200um diameter) connected to a syringe containing drug solution of choice. For IC administration, a postauricular skin incision to expose muscle overlying the temporal bone which was then separated and retracted to expose the bony wall of the posterior semicircular canal. A small hole was made using the tip of a myringotomy blade (Beaver-Visitec, Waltham, MA) to accommodate a customized polypropylene tube (OD: 100-200um) connected to a 10ul Hamilton syringe. The plastic tubing in both IB and IC administration were immobilized and sealed in place using cyanoacrylate glue (Permabond, Pottstown, PA). Drugs for the IC route were prepared in mouse artificial perilymph (AP; in mM: 150 NaCl, 4KCl, 8Na2HPO4, 2NaH2PO4, 1.5 CaCl2, 1 MgCl2, and 10 glucose; pH 7.2-7.4) and loaded into the Hamilton syringe. The syringe was connected to the IC plastic tubing where 1-2ul of solution was slowly delivered into the perilymph over 30-60 sec (Supplementary. Fig.1).

**Figure 1:**
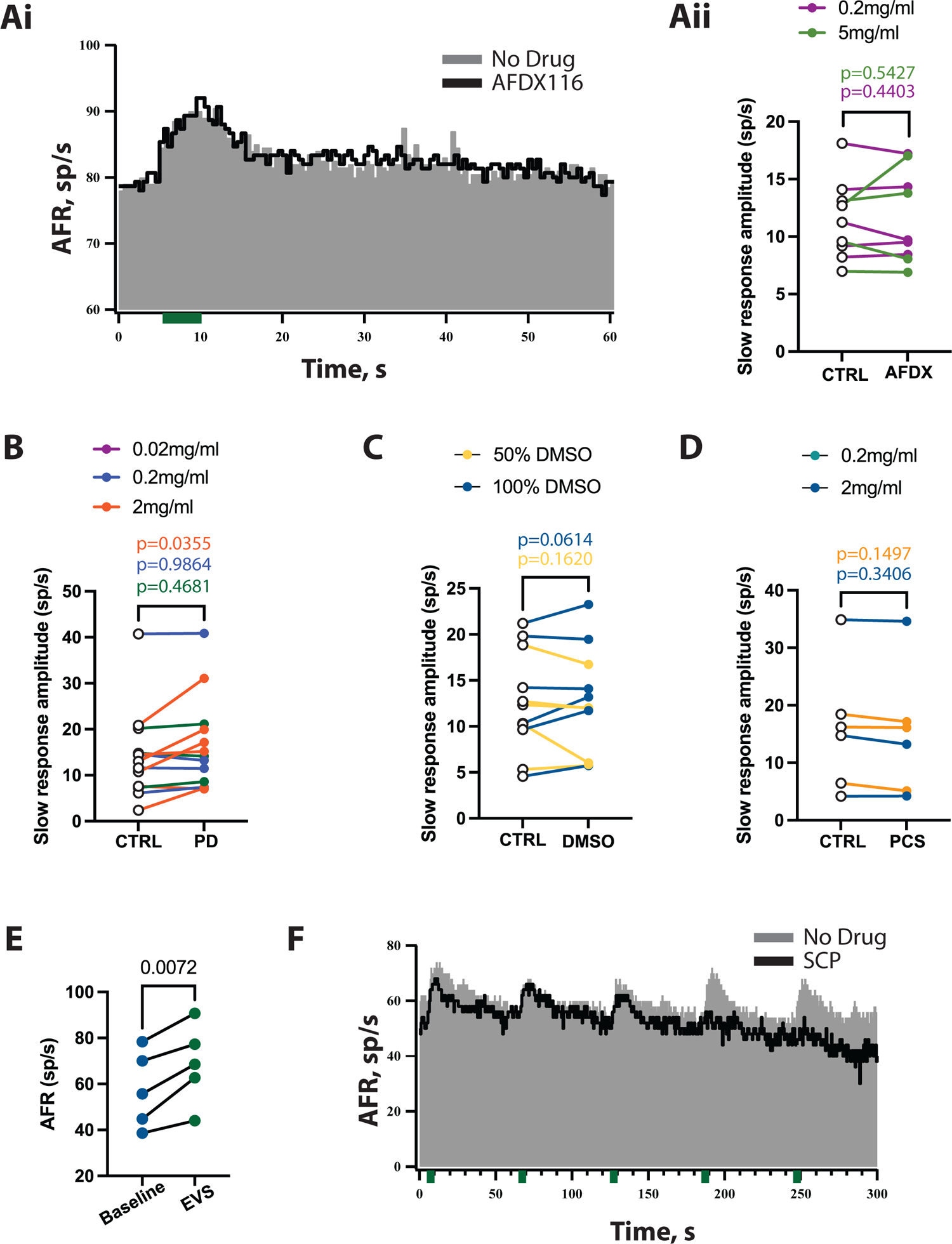
M2/M4 mAChRs do not drive EVS-mediated slow excitation. **Ai**, Representative average histogram showing EVS-mediated slow excitation in pre-(gray) and post-(black) IB administration of the M2mAChR antagonist, AF-DX116 at 0.2mg/ml. **Aii,** Mean amplitude of peak slow excitation before and after 0.2mg/ml (p=0.4403) and 5mg/ml (p=0.5427) AF-DX116 (AFDX). **B**, Mean amplitude of peak slow excitation before and after 0.02mg/ml (p=0.4681), 0.2mg/ml (p=0.9864) and 2mg/ml (p=0.0355) of the M4mAChR antagonist PD102807 (PD). **C,** Mean amplitude of peak slow excitation before and after IB administration of 50% (p=0.1620) and 100% (p=0.0614) DMSO in saline. **D,** Mean amplitude of peak slow excitation before and after IB administration of 0.2mg/ml (p=0.1497) and 2mg/ml (p=0.1497) of the M4mAChR antagonist, PCS 1055 (PCS). **E,** EVS stimulation in M2/M4 mAChR DKO mice lead to enhancement in firing rate of afferent neurons (n=5units/2 animals) (p=0.0072) **F**, Continuous response histogram showing EVS-mediated slow excitation during control conditions (CTRL, **grey)** and following blockade by the non-selective mAChR antagonist, scopolamine (**black**). (student’s paired t-test was used in all the above cases).

### Drug used

For this study, a number of drugs targeting mAChRs were used. Scopolamine, telenzepine (Yano et al., 2009; Ztaou et al., 2016), J104129 fumarate (Fedoce et al., 2016; Mitsuya et al., 1999), AF-DX116 (Kopf et al., 1998; Micheletti et al., 1987), PD102807 (Augelli-Szafran et al., 1998), VU0255035 (Sheffler et al., 2009), 4-DAMP (Andersson et al., 2011), DAU 5884 hydrochloride (Doods et al., 1993; Lorincz et al., 2008) and VU0238429 (Bridges et al., 2009; Foster et al., 2014) were all obtained from Bio-techne/Tocris (Minneapolis, MN). PCS1055 (Croy et al., 2016) and ML375 (Berizzi et al., 2018; McGowan et al., 2017; Radu et al., 2017) were obtained from Sigma Aldrich (St. Louis, MO). Stock solutions for scopolamine, telenzepine and J104129 were prepared in water and diluted to their final working solutions in saline for IP administration or in artificial perilymph for IC administration. IP doses are reported as mg/kg while IB and IC doses are reported as mg/ml. Stock solutions of AF-DX116, PD102807, PCS1055, VU0255035, 4-DAMP, DAU 5884 hydrochloride, VU0238429, and ML375 were prepared in 100% DMSO per manufacturer’s instructions. Working solutions for AF-DX116, PD102807, PCS1055, VU0238429, DAU 5884, and VU0255035 were prepared in DMSO/vehicle (where vehicle was saline or AP) at various dilutions: PCS1055 (0.2 mg/ml) at 1%; AF-DX116 (0.2 mg/ml), PD102807 (0.02 and 0.2mg/ml), PCS1055 (2 mg/ml), and VU0238429 (0.02 mg/ml) at 10%; VU0238429 (0.2 mg/ml) at 50%; while AF-DX116 (5 mg/ml) and PD102807 (2 mg/ml) remained in 100% DMSO.

Previous work demonstrated that IP administration of DMSO at volume less that 100ul had minimal effects of vestibular-evoked sensory potentials (Lee et al., 2017), therefore IP administration of any working drug solution in 100% DMSO was provided at a volume of 25ul DMSO for every 10g of the mouse’s weight.

The selection of a particular route of administration was governed by a number of often interrelated experimental constraints including the predictions about a particular drug’s access to the inner ear, procedural complexity of the administration route (IC>IB>IP), drug solubility requirements, and avoiding or minimizing undesired, off-target effects of drug and/or solvent (e.g., DMSO). For drugs that were readily soluble in physiological saline (e.g., scopolamine, telenzepine, and J104129), we would first start with IP injections and assess their efficacy in blocking mAChRs driving EVS-mediated slow excitation. Previous studies were consulted to predict dosing strengths. The likelihood an administered drug might access the inner ear was assessed using the SwissADME website that relies on the physiochemical and pharmacokinetic properties of each drug to make predictions how that drug moves across biological membranes (Daina et al., 2017; Daina & Zoete, 2016). While the SwissADME model emphasizes absorption in the gut and movement across the blood brain barrier (BBB) to enter the CNS, it has been used by our group and others to have potential insight about drug entry into the inner ear (Lee et al., 2017; Salt et al., 2019; Walia et al., 2021). This is best captured in the SwissADME egg plot where drugs readily absorbed from the gut are found within the egg white and drugs that also cross the BBB into the CNS are featured in the egg yolk. We generated a boiled egg plot for the full list of drugs employed in this study (Supplementary. Fig. 1). Six drugs would be predicted to cross the BBB, while five would be freely absorbed from the gut. VU0255035 is the only drug outside the egg white and would be predicted to do neither.

### VsEP

The linear VsEP is a compound action potential elicited from otolithic vestibular afferents, originating from the saccule and utricle, as a population response to transient linear acceleration and recorded on the surface of the scalp. The first three components of the VsEP (P1, N1 and P2) were scored and analyzed. Animal preparation and VsEP recording procedures were done in accordance with previously published works by Jones and colleagues for the mouse (Jones et al., 1999; Jones et al., 2002; Jones & Jones, 1999; Lee & Jones, 2018). Briefly, animals were anesthetized using IP urethane/xylazine (1.6g/kg, 20mg/kg). Animals were placed on a heating pad in a supine position and maintained at 37°C using a rectal thermistor and closed-loop monitoring system. Recording electrodes were placed epidurally on the nuchal crest (aka non-inverting electrode), behind the right pinna (i.e., inverting electrode), and on the hip (i.e., ground). The head was then secured to a voltage-controlled mechanical shaker via a custom-made head mount. Linear acceleration pulses were generated by using custom software. A linear voltage ramp of 2-ms duration was applied to an electromagnetic shaker (model ET-132-203; LabWorks, Costa Mesa, CA). Stimuli were presented at a rate of 17 pulses/s. Stimulus amplitude was calibrated and monitored using an accelerometer mounted on the shaker platform. Stimulus amplitude was quantified in decibels relative to the reference unit of jerk (specifically dB re: 1.0 g/ms, where 1.0 g=9.8 m/s^2^), and ranged from −15 to +6 dB re: 1.0 g/ms. To eliminate any potential auditory contributions to the VsEP data, vestibular stimuli were presented with a binaural forward masker of broadband noise (50-50,000 Hz bandwidth, 90dB SPL) via a free-field speaker within the recording booth. The resulting electrophysiological signal was amplified 200,000-fold using a Grass P511 amplifier and filtered @ 0.3-3 kHz. The signal was digitized at the onset of each stimulus (1,024 points at 10us/point). Signal averaging was employed to enhance the signal-to-noise ratio and generate a mean VsEP response trace. To do this, separate average traces of each polarity were first collected (128 sweeps in each direction) and then subsequently summed to generate a final VsEP trace from a total of 256 sweeps. After acquiring baseline VsEP metrics, the mAChR agonist oxotremorine-M was administered intraperitoneally at a concentration of 2 mM (10μl/g of animal). We then recorded VsEPs at +6 dB (re: 1.0 g/ms) at 2.5-minute intervals and assessed VsEP threshold every 10-15 minutes for a duration of 20-30 mins.

### Behavioral experiments

Mice aged 10 to 14-weeks of both sexes were used for all the behavioral experiments. The same animal was used for various experiments and was tracked using an animal ID microchip (Manruta, 1.25×7mm). The microchip was injected under the skin after anesthetizing the mouse with isoflurane. Mice were allowed to recover for 3 days before start of any of the behavioral experiments.

#### Open field activity

To test motor and exploratory behavior, open field activity was monitored. Mice were brought to the testing room at least 1hrs before start of the test for acclimation to the room and lighting. Mice were then placed in the center of a well-illuminated circular arena measuring 17-inches in diameter. The animal’s activity was recorded using an overhead camera (Basler acA1920-155um) and video data were captured for post experimental analysis. For open field activity in the dark, the arena was illuminated with infrared lights (Axton, Smart-B series, model AT-8S-B, 850nm). Mice were acclimated from low light to dark conditions for at least 1-hr before recording. The room with arena was illuminated with red light between the experiments for experimenter to visualize the area. Analyses were done using behavioral analysis software, EthoVision XT (version 15, Noldus Information Technology). The arena was wiped with 70% ethanol between animals and allowed to dry before beginning the next set of observations.

#### Balance beam

Motor coordination and balance were tested using a custom balance beam assay set up in the lab. For these experiments, the balance beam consisted of an elevated aluminum bar measuring 80cm long and available in two widths (9 or 18 mm: testing vs training beam). An enclosed box containing nesting materials and a red enrichment house from their home cage was located on one end of the beam and served as a target location for animals placed on the starting position found on the opposite end of the beam. The starting position was well illuminated with 60W light and the enclosed box was darker to encourage animals to traverse the beam. On the first two days, mice were trained on the balance beam for a total of 3 trials per day. On Day 3, mice were placed on the smaller beam and the mean time needed to reach the home box was computed from two trials. All beam sessions were recorded using an integrated camera in a Dell Inspiron 15 laptop and further analyzed using the behavioral analysis software EthoVision XT (Version 15, Noldus Information Technology). The beam and home cage were wiped with 70% ethanol and allowed to dry before and after each animal.

#### Gait analysis

To test motor function, locomotion, and coordination, a gait analysis was performed within the URMC Behavioral Sciences Facility Core. For recording gait, mice were placed on a walkway consisting of a transparent plexiglass lane (enclosed on all four sides). A high-speed video camera (100 frames/s) was mounted underneath the walkway permitting digital recordings of the underside of the mouse including the soles of their feet. Mice were trained to walk from a start position along the walkway to a goal box with enclosed housing located on the opposite end. After three training runs were performed for each animal to teach the subject to move toward the goal box, the animal was then recorded walking across the walkway anywhere from 3 to 10 times (depending on the quality of the digital capture of the subject’s gait). In-house analysis software was used to identify initial foot contact, stance duration, stride length, stride duration, foot liftoff, swing duration, track width, and toe spread data for each foot. A single session video was used to train the software based on its representation of normal mouse locomotion. For each session, video was previewed to determine a minimum of four to six consecutive step cycles of consistent walking for video analysis. Step cycles in which animals drifted to the sides or explore the chamber were excluded from analyses. The surface of the plexiglass walkway was cleaned with 70% ethanol between each session.

#### Postural sway and Center of pressure (COP) measurements

To measure postural sway, a low load miniature force platform (HE6X6, AMTI Watertown, MA) was used. The platform measured 6×6 inches. The set-up was walled off by a four-sided plexiglass enclosure with 3 black opaque and one transparent side to visualize the animals. The enclosure was free standing and did not contact or obstruct the force plate. Mice were brought to the vestibular behavioral core at least 30 mins before the start of experiment. After room acclimation, each mouse to be tested was first placed in an enclosed box of similar dimensions to the recording set-up for 5 mins for further acclimation.

Afterwards, the mouse was transferred to the enclosed force plate and the animal’s weight was obtained. The mouse was allowed to explore the platform for 1-2 min before starting postural measurements. We then acquired 30, 1-sec sampling blocks of the three-dimensional forces and moments associated with the animal’s standing on the force platform. Each sample was only acquired when the mouse was down on all four limbs and not rearing or grooming. Each 1-sec sampling block consisted of 300 samples, each of which can be used to compute the animal’s downward force at each of the 300 time points. Collectively, when plotted, the (x, y) co-ordinates of these 300 samples form an ellipse that aligns with the postural sway space of that animal along the fore-aft axis. The data is fit with a 95% confidence ellipse and the area of this ellipse was calculated and used to indicate the amount of sway exhibited by a mouse on a particular trial. For each ellipse, a single center of pressure (COP) value is had by determining the point of intersection between the long and short axis of the ellipse. After obtaining baseline COP values, mice are then transferred to an enclosure on orbital shaker for the vestibular challenge where they are rotated at 125rpm (4cm displacement) for 5-mins. Afterwards, mice were then immediately transferred back to force plate where further COP measurements were taken at t=0, 1, 3, 5, 10, and 15min. All enclosures were cleaned with 70% ethanol between animal test sessions.

#### Thermal recordings

Thermoregulatory responses of mice to vestibular challenge were recorded using an infrared camera (FLIR-E60, Flir Systems, Wilsonville, OR). Animals were brought to the vestibular behavior core at least 30 mins before the start of the experiment. Animals were placed in a 2×4 inch plexiglass container on the orbital shaker with the infrared camera mounted above to record mouse body temperature in real time. Camera was mounted on the orbital shaker so that it rotated in sync with mouse container. Mice were allowed to explore the container 5 minutes before the start of the experiment. Baseline temperatures were recorded for 5 mins in the absence of rotation. After baseline, mice were rotated at 75rpm on the orbital shaker for 20-mins while the animal’s temperature was continuously monitored. After rotation, temperature recordings were made for an additional 20 minutes. All temperature data were analyzed using FLIR software. For our studies, we specifically analyzed head and tail temperatures. Head temperature was recorded at the mid-point between the two ears while tail temperature was taken at 1/3^rd^ of tail length from the body. Head and tail temperatures were assessed in two-minute intervals. The chamber where the animals were rotated was cleaned between test sessions (Tu et al., 2017).

#### Forced Swim Test

The forced swim testing was carried out in a 5L glass flask filled halfway with water maintained at room temperature at 25 degree C. Mice were placed in the water for a maximum period of 6 min without the opportunity to escape. At the end of the 6 min, mice were removed, dried, and placed back into their home cage. Video of behavior during swim test and post-swim recovery in the home cage were recorded for later scoring and evaluation. Water was changed between animals. The analysis of the forced swim test was divided into 2 parts: 2-min training period and 4-min of testing period. Animal was scored on ‘Freeze’ and ‘Swim’ behavior. Forced swim test was conducted and analyzed by the mouse Behavioral Sciences Facility Core URMC and the Holt Lab.

#### Post-Forced Swim Test

Animal was placed in a cage post swim test to record their post-swim behavior. The cage was kept warm using a heat lamp. Animals were observed for instability in their gate and posture for 10mins.

### Statistical procedures

The effects of different pharmacological treatments on afferent discharge rate and EVS-mediated slow excitation were assessed using a paired t-test. Data were tested for normality before using any statistical procedure. One way ANOVA with Tukey’s multiple comparison was used to: (1) Compare the effects of multiple drugs on the same unit; (2) Compare EVS-mediated slow and fast excitation; and (3) VsEP amplitude and latency among different genotypes. rmMANOVA with post hoc Bonferroni was used to analyze effect of time or stimulus intensity on, 1) VsEP amplitude and latency as well as, 2) Thermoregulation post VC, between control, M3 mAChR het and M3 mAChR KO mice. All statistical analyses were done in Graph Pad-Prism (GraphPad). Values, expressed as means, SEM, and outcome parameters including p-values, F-statistics, t-statistics, and effect sizes (Cohen’s d) are reported in the text and/or figures.

## Results

### M2/M4 mAChRs do not mediate EVS-mediated slow excitation

Previous studies in several vertebrate species have revealed that EVS-mediated slow excitation is driven by mAChR activation (Holt et al., 2017; Lee et al., 2021; Schneider et al., 2021). Additional electrophysiological and pharmacological data indicate that the resulting slow excitation likely utilizes an M-current mechanism, whereby activation of mAChRs results in the closure of KCNQ channels leading to an increase in afferent impedance and excitability (Holt et al., 2017; Perez et al., 2009; Ramakrishna et al., 2021). While the mAChR subtypes (M1-M5) driving EVS-mediated slow excitation have not been identified, an M-current mechanism would implicate the odd-numbered, but not even numbered, mAChRs on the basis of G-protein coupling and downstream signaling cascades (Brown & Adams, 1980; Lerche et al., 2000; Schroeder et al., 2000; Søgaard et al., 2006; Suh & Hille, 2002; Suh et al., 2006; Wang et al., 1998).

To evaluate if even-numbered (M2/M4) mAChRs played any role in EVS-mediated slow excitation, we acquired extracellular recordings of EVS-mediated slow excitation from mouse vestibular afferents before and after the administration of M2- or M4-selective mAChR antagonists. Figure 2.1Ai shows a typical EVS-mediated slow excitatory response in a regular vestibular afferent before and after the intrabullar (IB) administration of the M2-selective antagonist AF-DX116 (30 μl @ 5 mg/ml). Under control conditions (gray), the average response histogram reveals a slow excitation which peaks at 10 spikes/s and persists for over 30 seconds. At 10-30mins (10 minutes in this example) following the delivery of AF-DX116 into the middle ear space, EVS-mediated slow excitation remained relatively unchanged. At two concentrations (0.2 and 5 mg/ml), AF-DX116 had no consistent effect on EVS-mediated slow excitation in a total of nine afferents (Fig. 1Aii). The mean peak amplitude of EVS-mediated slow excitation under control conditions was not significantly different from those obtained in 0.2 mg/ml AF-DX116 (n=5, 12.16± 1.797 vs 11.84± 1.681 spikes/s, t(4)=0.8559, p=0.4403, student’s paired t-test) or 5 mg/ml AF-DX116 (n=4, 10.58± 1.442 vs 11.44± 2.395 spikes/s, t(3)=0.6846, p=0.5427, paired t-test). These data suggest that activation of M2mAChRs does not contribute to EVS-mediated slow excitation of mouse vestibular afferents.

**Figure 2:**
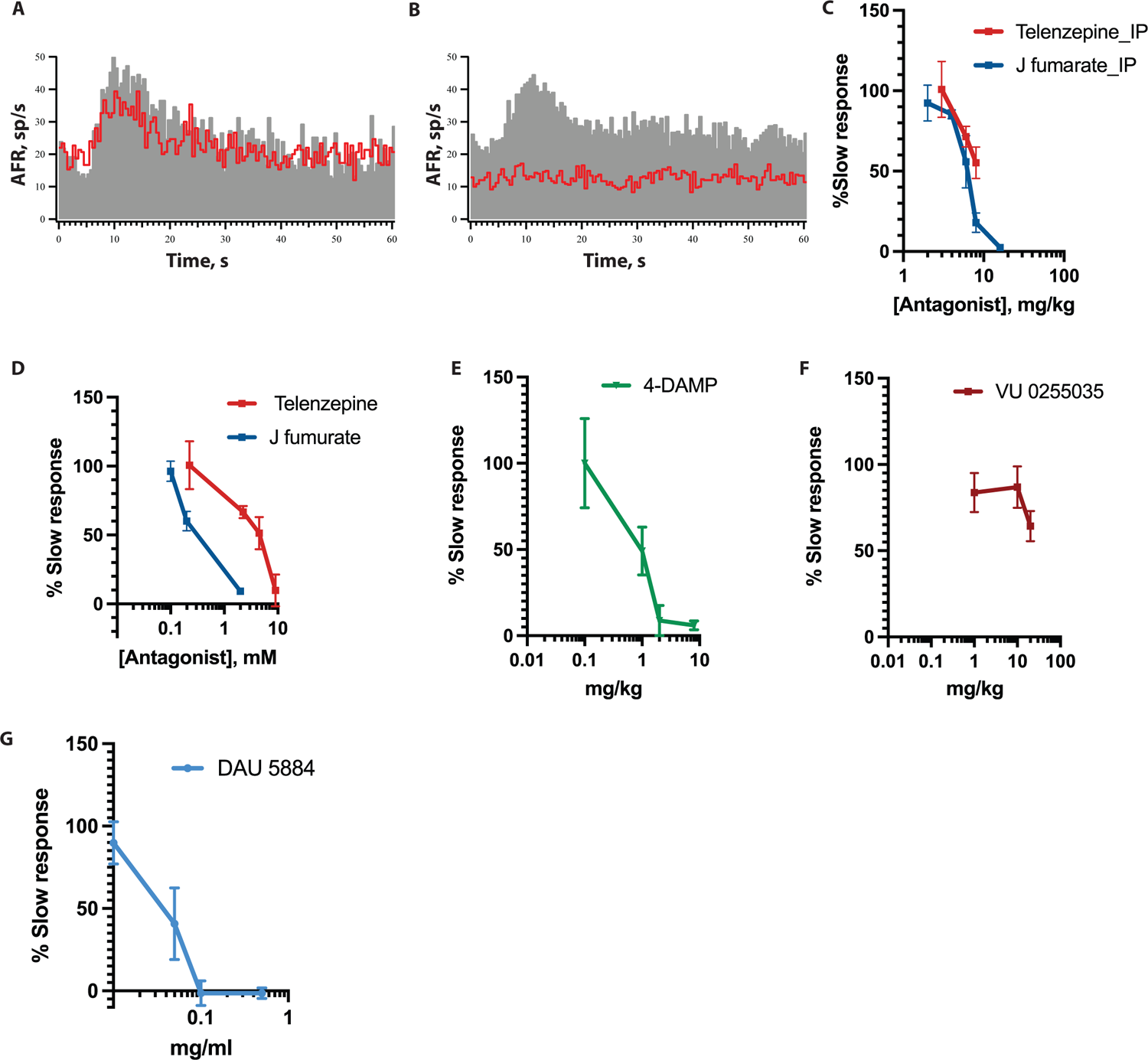
M3mAChR antagonists substantially reduce slow excitation compared to M1mAChR antagonists. **A, B**, Average response histogram showing EVS-mediated slow excitation before and after the intraperitoneal (IP) administration of the (A) M1mAChR antagonist, telenzepine dihydrochloride and (B) M3mAChR antagonist, J104129 fumarate, both at 8 mg/kg. **C,** Dose response curve showing % EVS-mediated slow excitation remaining after IP administration telenzepine and J104129. **D,** Direct administration of telenzepine and J104129 fumarate into perilymph via intracanal (IC) administration improves accessibility to afferent synapses in the sensory neuroepithelia. IC dose response curve for both drugs are shown. **E,** IP dose-response curves for the M3mAChR selective antagonist, 4-DAMP, and M1mAChR selective antagonist, VU 0255035, are shown. **H**. Dose response curve for IB administration of M3 mAChR antagonist, DAU 5884. (n=3-5 animals/drug/concentration).

We next measured EVS-mediated slow excitation in a total of 12 afferents before and after the IB administration of the selective M4mAChR antagonist PD102807 at three different concentrations (0.02, 0.2, 2 mg/ml). Neither concentration proved to be an effective blocker (Fig. 1B). As compared to their controls, EVS-mediated slow excitation was not significantly different at 0.02 mg/ml (14.1± 3.72 vs 14.62±3.62 spikes/s, t(2)=0.8883, p=0.4681, student’s paired t-test) or at 0.2 mg/ml (18.25±7.69 vs 18.24 ± 7.63 spikes/s, t(3)=0.0184, p=0.9864, student’s paired t-test). Surprisingly, peak EVS-mediated slow excitation amplitude in 2mg/ml PD102807 was significantly greater than control (11.52 ± 2.56 vs 16.26 ± 3.659 spikes/s, t(5)=2.859, p=0.035, student’s paired t-test). While 0.02 and 0.2 mg/ml PD102807 were dissolved in 10% DMSO, 2mg/ml required 100% DMSO to maintain solubility. We suspect that the significant increase in EVS-mediated excitation during the 2mg/ml concentration may be a product of DMSO and/or high concentration of PD102807. As additional controls, afferent firing background discharge rates and EVS-mediated slow excitation were measured before and after the IB administration of 50% or 100% DMSO alone. While 100% DMSO appeared to increase the amplitude of EVS-mediated excitation in several units (Fig. 1C), slow response amplitudes in 50 or 100% DMSO were not significantly different from controls (50% DMSO: 11.9 ± 2.18 vs 10.5 ± 2.07 spikes/s, t(4)=1.712, p=0.1620, student’s paired t-test; 100% DMSO: 13.3 ± 2.6 vs 14.57 ± 2.5 spikes/s, t (5)=2.405, p=0.0614, student’s paired t-test). Furthermore, neither concentration of DMSO significantly affected background discharge rates (50% DMSO: 40.56 ± 13.48 vs 44.53 ± 12.2 spikes/s, t(4)=1.323, p=0.59, student’s paired t-test; 100% DMSO: 46.09 ± 14.08 vs 51.17 ± 13.23 spikes/s, t(5)=1.859, p=0.35, student’s paired t-test (data not shown)). These data suggested that either these are combined, nonspecific effects of DMSO and PD102807, or that possibly that blockade of M4mAChRs somehow gives rise to an enhanced EVS-mediated slow excitation. To further test if the enhancement seen in 2mg/ml PD102807 was due to selective blockade of M4mAChRs, we used another selective M4mAChR antagonist, PCS1055 (Fig 1D). However, PCS1055 did not show significant enhancement nor blockade of EVS-mediated slow excitation at two different concentrations (0.2mg/ml: 13.71 ± 3.68 vs 12.79 ± 3.85 spikes/s, t(2)=2.285, p=0.1497, student’s paired t-test; 2mg/ml: 17.94 ± 9.0 vs 17.35 ± 9.02 spikes/s, t(2)=1.241, p=0.3406, student’s paired t-test). Similar to our observations with the M2mAChR antagonist, these data suggest that M4mAChRs do not play a role in EVS-mediated slow excitation.

One last set of experiments aligns quite well with our pharmacological observations using the M2/M4 mAChR antagonists. In our pursuits to characterize the mAChR subtypes underlying EVS-mediated excitation, we also performed afferent recordings from M2/M4-mAChR double-knockout animals during efferent stimulation as exemplified by the continuous histogram shown in Figure 1E where the each one of the five EVS-shock trains results in a typical slow excitation. In five afferent recordings from two mice, EVS-mediated slow excitation (No EVS stimulation vs EVS stimulation: 57.50± 7.46 vs 68.63±7.75, t(4)=5.063, p=0.0072, student’s paired t-test) was observed and appeared quantitatively similar to that seen in our control animals. That the EVS-mediated slow excitation in these animals was attributed to the activation of mAChRs was confirmed by blockade with the non-selective mAChR antagonist scopolamine as previously reported in C57Bl/6 animals (Fig 1F) (Lee et al., 2021; Schneider et al., 2021). Collectively, these data are consistent with the prediction that odd, but not even-numbered, mAChRs are driving EVS-mediated excitation in mice.

### M3mAChR antagonists substantially reduce slow excitation as compared to M1mAChR antagonists

To probe the role for odd-numbered mAChRs in EVS-mediated slow excitation, we obtained extracellular afferent recordings in C57Bl/6 mice during EVS stimulation before and after the application of selective M1, M3, and M5mAChR drugs. First, we determined the relative sensitivity of EVS-mediated slow excitation to the IP administration of the M1mAChR antagonist, telenzepine, or M3mAChR antagonist, J104129 fumarate (J104129), at a range of increasing concentrations (Fig. 2A-D). During IP administration of both drugs at similar concentrations, J104129 was generally more effective at blocking EVS-mediated slow excitation compared to telenzepine. The relative differences between these two drugs are exemplified in Figure 2A and 2B where average response histograms show EVS-mediated slow excitation before and after the IP administration of telenzepine and J104129, both at 8mg/kg, respectively. In these examples, J104129 is a more potent blocker than telenzepine. In agreement with these individual examples, superimposition of the inhibitory dose response curves for the IP administration of the two mAChR antagonists reveal that a comparable block of EVS-mediated slow excitation required lower doses of J104129 (Fig. 2C). There is a greater than two-fold difference in the dose needed to block 50% of the response (J104129 f vs Telenzepine: ID50 (inhibitory dosage): 6.03 vs 14.43 mg/kg).

While IP administration of telenzepine and J104129 clearly reached the ear to block EVS-mediated slow excitation, it required concentrations 2-4 times higher than those used to target mAChRs in other studies (Mitsuya et al., 1999; Yano et al., 2009; Ztaou et al., 2016), suggesting that higher systemic doses of telenzepine and J104129 are needed in order to reach the appropriate blocking concentrations in the perilymph. From our previous work (Lee et al., 2021), we have identified that (1) the rules governing drug entry into the inner ear may be at odds with entry of that same substance into the CNS; and (2) not all substances, administered through the IP route, readily enter the inner ear in the same way. We submitted a number of mAChR antagonists using the SwissADME portal (http://www.swissadme.ch) to determine lipid solubility (WLOGP) vs polar surface (TPSA) for each drug (Supplementary Fig. 1). The drugs present in egg ‘yolk’ can cross the blood-brain barrier (BBB) while drugs present in the egg white plot can be readily distributed gut following absorption but cannot readily cross the BBB (Daina & Zoete, 2016). Based on the boiled egg plot, both telenzepine and J104129 fumarate are predicted to not enter the CNS readily, and presumably as such entry into the inner ear may also require substantially higher doses. To address this problem and to ensure that these two drugs had access to vestibular efferent synapses, we directly administered telenzepine and J104129 into perilymph through a intracanal (IC) administration (Lee et al., 2021). As with IP administration, IC administration of J104129 fumarate was again more effective at blocking EVS-mediated slow excitation compared to IC telenzepine (IC50: 0.17mM vs 3.12 mM, Fig 2.2F). While these data suggest a major role for M3mAChRs, it may also be proposed that M1mAChRs play a minor role. Since the dose response curves overlap, telenzepine and J104129 may also begin to block M3 and M1mAChRs, respectively at higher concentrations.

To further probe if M3 and/or M1mAChRs are involved in EVS-mediated slow excitation, we next characterized EVS-mediated slow excitation in the presence of several other M1 and M3mAChR antagonists. In dose-response curves (Fig. 2E-F), the effects of increasing doses of the M3mAChR antagonist 4-DAMP (ID50: 0.9655mg/kg) and M1mAChR antagonist VU0255035 (ID50:> 20 mg/kg) on EVS-mediated excitation are shown. Both 4-DAMP and VU0255035 were administered through the IP route as these drugs were previously demonstrated to enter the CNS through systemic administration (Andersson et al., 2011; Rahman et al., 2020; Sheffler et al., 2009), and we reasoned that they would likely enter the ear as well. Again, mirroring the data with telenzepine and J104129, the M3mAChR antagonist 4-DAMP was a much more effective blocker of EVS-mediated slow excitation than VU0255035. It should be noted, however, that VU0255035 does exhibit some block at the lowest concentrations used suggesting the M1mAChRs may again play a small role. Additionally, we administered VU0255035 via IC route as the boiled egg plot suggested that the VU 0255035 might not readily enter the inner ear though previous work suggested that VU0255035 do enter CNS (see methods). When used at 0.2ug/μl, the maximum concentration that can be dissolved in artificial perilymph, we saw a 41% reduction in EVS-mediated slow excitation (data not shown). Finally, we also administered the M3mAChR antagonist DAU 5884. Though the Swiss ADME site indicated that DAU5884 should access the CNS (and ear) after systemic administration, our initial recordings with IP administration were associated with a number of off-target affects that impacted afferent excitability and made interpretation difficult. We then elected to administer DAU5884 using the IB route which mitigated these issues. The dose-response curve for the effects of DAU5884 revealed that it is quite potent at blocking EVS-mediated slow excitation (Fig. 2G). Hence, in all the cases, M3mAChR selective antagonists substantially reduced the EVS-mediated slow excitation compared to M1mAChR selective antagonists.

### Activation of M5mAChRs enhance the background firing rate of afferent neurons but does not alter EVS-mediated slow excitation

Prior studies have demonstrated that all multiple mAChR subtypes are present in the vestibular end organs of guinea pigs, rat, and humans (Ishiyama et al., 1997; Li et al., 2007; Yao et al., 2011; Wackym et al., 1996). Thus far, pharmacological data and the presence of EVS-mediated slow excitation in M2/M4 double-KO mice suggest that M2 and M4mAChRs are not involved, while other pharmacological data implicate a major role for M3mAChRs and perhaps a minor role for M1mAChRs. In order to explore a remaining role for M5mAChRs, we sought to characterize if EVS-mediated slow excitation could be modified by drugs known to selectively modulate M5mAChRs. As there is a lack of commercially-available M5mAChR-selective antagonists, we used the M5mAChR positive allosteric modulator (PAM), VU0238429. If M5mAChRs contribute to EVS-mediated slow excitation, VU0238429 would be predicted to enhance the amplitude of the EVS-mediated slow response. For this set of experiments, EVS-mediated slow excitation was measured before and after the IB administration of VU0238429 at 0.02 and 0.2 mg/ml. In Figure 3A, four trials of EVS-mediated slow excitation were acquired before 0.2 mg/ml VU0238429 was delivered to the middle ear space at t = 225s (blue box). EVS-mediated slow excitation changed very little over the next 17 minutes (slow response amplitude: 20.86 vs 16.25). In a total of 12 units, we found that IB administration of VU0238429 did not significantly enhance EVS-mediated slow responses either at 0.02mg/ml (14.92±2.51 vs 13.93±1.05 spikes/s, t(3)=0.4174, p=0.7044, student’s paired t-test) or 0.2mg/ml (12.58± 1.81 vs 10.81±1.86, n=9, p=0.1478, Tukey’s multiple comparison; Fig. 3B). However, subsequent application of the non-selective mAChR antagonist scopolamine, following IB VU0238429, was able to completely block EVS-mediated slow excitation in seven units in the 0.2 mg/ml group (12.58±1.81 vs −0.34±0.71, n=7, p=0.0056, Tukey’s multiple comparison) (2 units were lost before assessing scopolamine’s effects).

**Figure 3.**
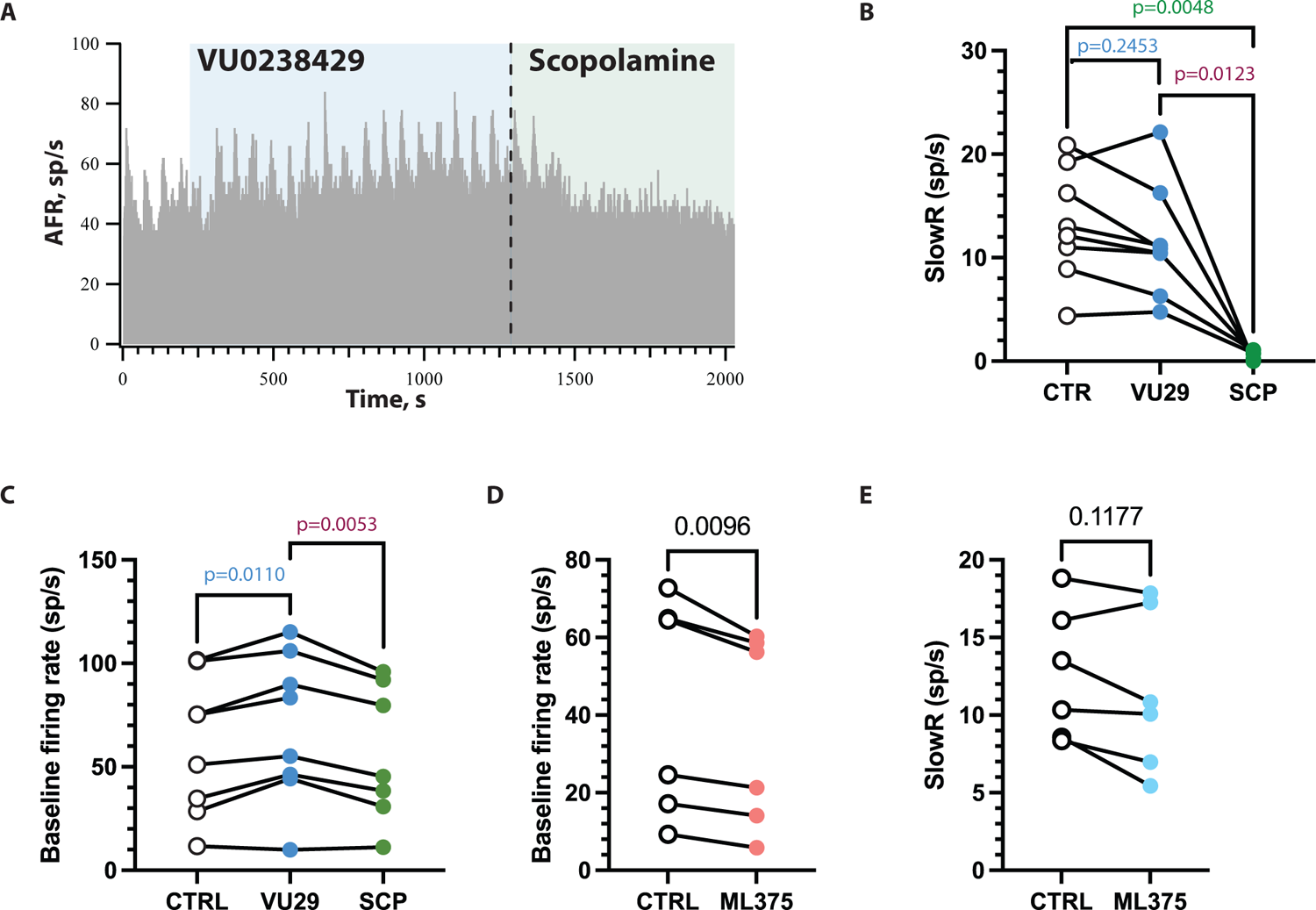
Activation of M5mAChRs enhances the background firing rate of afferent neurons but does not affect amplitude of EVS-mediated slow excitation. **A.** Representative continuous histogram showing background firing rate and EVS-mediated slow excitation during CTRL (white box), in presence of M5mAChR PAM VU 0238429 (blue box), followed by IP scopolamine (green box). **B,** Peak EVS-mediated slow excitation amplitudes during pre-drug (CTRL), post-IB VU0238429 (VU29, 0.2 mg/ml), and post IP Scopolamine (SCP, 2 mg/kg). **C**, Baseline afferent firing rate of afferent during pre-drug (CTRL), post-IB VU0238429 (VU29, 0.2 mg/ml), and post-IP Scopolamine (SCP, 2mg/kg). **D,** Baseline afferent firing rates of afferents pre-drug (CTRL) and post-IP administration of the M5mAChR NAM, ML375 (5 mg/kg). **E**, EVS-mediated slow excitation amplitude of afferents pre-drug (CTRL) and post-IP administration of ML375 (5 mg/kg).

Interestingly, despite having no effect on EVS-mediated slow excitation, we observed an enhancement in the background firing rate of afferents during treatment with VU0238429 (Fig. 3A, C). In the example shown in Figure 3A, the baseline firing rate hovers near 40 spikes/s in the beginning but gradually increases to about 55 spikes/s over the course of IB VU0238429. An enhancement in the baseline was noted at both the 0.02mg/ml concentration (24.65±14.41 vs 31.23±15.77, n=4/4, t(3)=3.409, p=0.0422, student’s paired t-test) and the 0.2mg/ml dose (57.67±10.72 vs 65.43±11.64, n=9, p=0.0184Tukey’s multiple comparison). That the VU0238429-mediated slow enhancement of baseline discharge was attributed to mAChRs was confirmed by its ablation during the subsequent IP administration of scopolamine (Fig 3A, 3C). Baseline firing rates fell back to pre-VU0238429 levels (VU0238429 vs scopolamine: 0.2mg/ml: 65.43±11.64 vs 56.18±12.47, p=0.0166). These pharmacological data suggest that the activation of M5mAChRs can enhance afferent firing independent of the mAChRs driving EVS-mediated slow excitation and over a longer (slower) time scale. To provide further evidence for M5mAChRs, we also employed the M5mAChR negative allosteric modulator (NAM), ML375. Opposite to our observations with the VU0238429, IB ML375 consistently reduced the baseline firing rate of mouse vestibular afferents (Fig. 3D) (42.21±11.50 vs 36.03±10.20, t(5)=4.070, p=0.0096, student’s paired t-test), but like VU0238429, ML375 did not result in consistent effects on EVS-mediated slow excitation (Fig. 3E) (12.62±1.743 vs 11.41±2.109, t(5)=1.888, p=0.1177, student’s t-test). These data suggest that M5mAChRs are also activated by EVS stimulation to give rise to a much slower afferent excitation that appears to work through a mechanism different than that used by M1/M3 mAChRs described in earlier figures.

### Characterization of resting discharge rate, and EVS mediated fast and slow excitation in C57BL/6J wild type (WT), Heterozygous M3 mAChR (M3 Het) and M3 mAChR KO (M3 KO) mice

Based on the aforementioned pharmacological evidence, we hypothesized that activation of M3mAChRs underlie EVS-mediated slow excitation in mouse vestibular afferents, while M1mAChRs may play a smaller role. To further test this assertion, we sought to characterize EVS-mediated slow excitation in M3mAChR knockout mice (M3KO), with the expectation that vestibular afferents in these mice will have attenuated or missing EVS-mediated slow responses. We compared electrophysiological data from M3KO animals to C57Bl/6 controls and heterozygous animals (M3Het). In this set of experiments, tribromoethanol was used for anesthesia (see Methods) as we have pharmacological evidence that urethane anesthesia attenuates EVS-mediated fast excitation while having no effect on EVS-mediated slow excitation (Lee C et al., ARO MWM (2021)). In the event that M3mAChR-KO animals were devoid of EVS-mediated slow excitation, we wanted to be sure that we had independent evidence that EVS neurons were still functionally viable and could be recruited with our standard electrical stimulation. EVS-mediated fast excitation, which is mediated by nAChRs, should remain intact and thus works well as an internal control, provided we avoid the use of urethane anesthesia.

We first characterized the resting discharge rates of vestibular afferents in control, M3Het, and M3KO animals. Using a normalized coefficient of variation (CV*) to describe the discharge regularity of vestibular afferents (Goldberg et al., 1984; Jones et al., 2008; Schneider et al., 2021), we classified vestibular afferents as regular (CV*<0.1) or irregular (CV*>0.1). For the purpose of this work, intermediate-firing afferents (0.1<CV*<0.2) were included in the irregular category. Resting discharge rates versus CV* among WT controls (blue, n=40 units/6 animals), M3Het (green, n=32 units/5 animals) and M3KO animals (red, n=60 units/6 animals) had similar distributions (Fig. 4A, top row) and were comparable to what has been previously reported in mammals (Goldberg & Fernandez, 1971) (squirrel monkey); (Jones et al., 2008) (mice); (Baird et al., 1988) (chinchilla). There were no obvious differences across the three groups. (Table 1).

**Figure 4.**
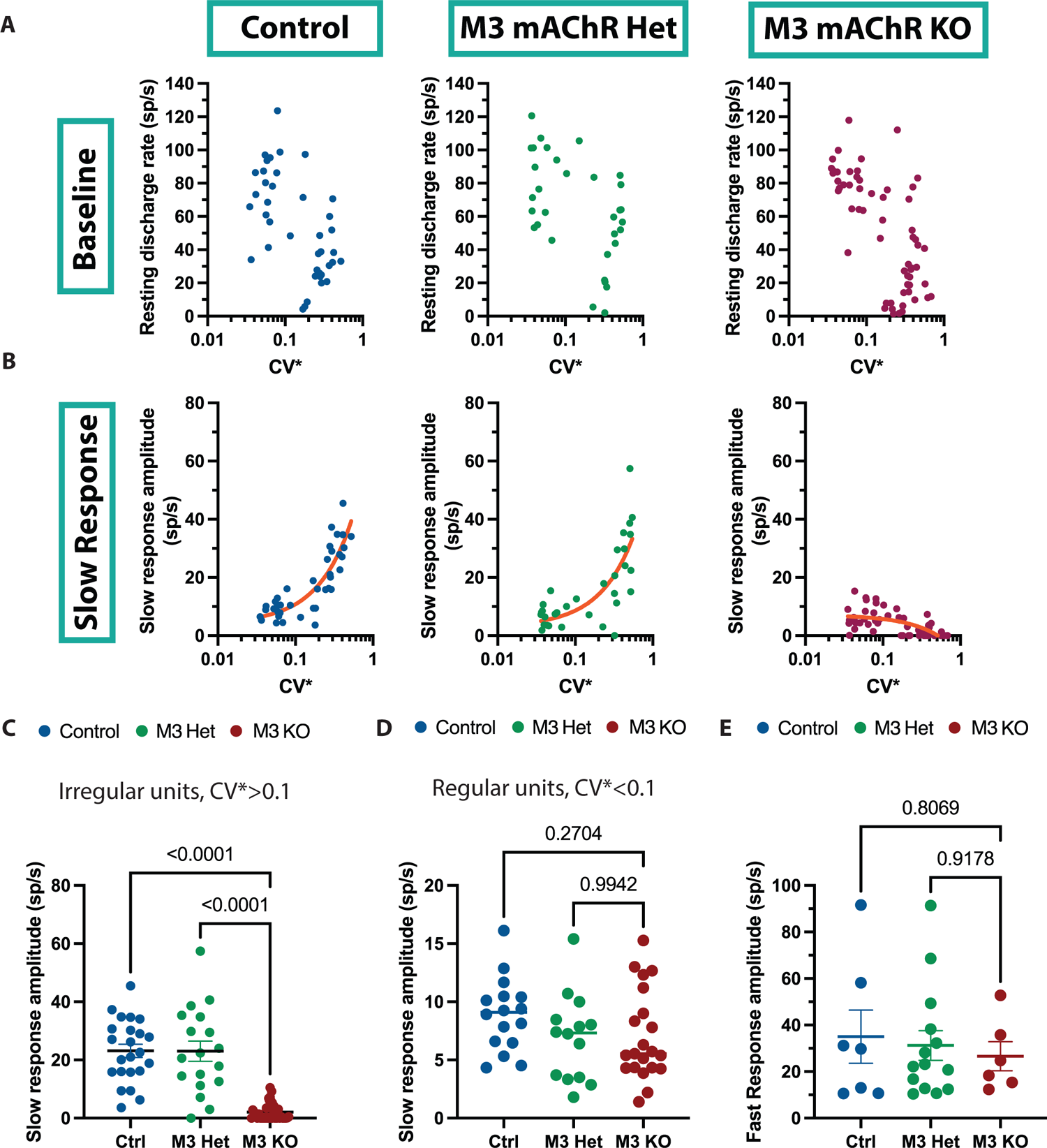
Characterization of resting discharge, EVS mediated slow excitation, and EVS-mediated slow excitation in C57BL/6J wild type (WT), M3Het (Chrm3^+/-^) and M3KO (Chrm3^-/-^) animals. **A, B**, Data for wildtype C57BL/6J mice (n=6, 40 units; Left Column, Blue); M3mAChR-Het mice (n=5, 32 units; Middle column, Green); and M3mAChR-KO mice (n=6, 60 units; Right Column) are shown. For each unit in each strain, **A** Resting discharge rates of vestibular afferents and **B,** peak slow response amplitude values, were plotted against CV*, a standardized measure of discharge regularity where low/high CV* values denote regular/irregular discharge, respectively. **Orange line** indicates linear fit of data. **C,** EVS-mediated slow response amplitude in irregular firing afferents (CV*>0.1) in wild type, M3 mAChR Het and M3 mAChR KO mice. **D,** EVS-mediated slow response amplitude in regular firing afferents (CV*<0.1) in wild type, M3 mAChR Het and M3 mAChR KO mice. **E,** Fast response amplitude greater than 10 spikes/s in all vestibular afferents from wild type, M3 mAChR Het and M3 mAChR KO mice are plotted.

**Table 1:** Mean resting discharge rate of primary vestibular afferents in Wild type, M3mAChR Het and M3mAChR KO mice.

|  | CTRL | M3 mAChR Het | M3 mAChR KO |
| --- | --- | --- | --- |
| Mean of resting discharge rate of all afferents | 54.31±4.79<br>(n=40) | 64.87±5.44<br>(n=32) | 51.02±4.323 (n=60) |
| Mean of resting discharge rate of regular firing afferents (CV*$<0.1$) | 78.10±5.39<br>(n=17) | 81.63±6.28<br>(n=14) | 81.62±3.32 (n=22) |
| Mean of resting discharge rate of irregular firing afferents (CV*$>0.1$) | 36.73±4.69<br>(n=23) | 51.83±7.05<br>(n=18) | 33.31±4.50 (n=38) |

We next characterized EVS-mediated slow excitation in control, M3Het, and M3KO animals and plotted mean response amplitudes versus CV* (Fig. 4B, bottom row). As previously reported (Brichta & Goldberg, 1996; Goldberg & Fernandez, 1980; Plotnik et al., 2002; Schneider et al., 2021), the amplitude of EVS-mediated slow excitation in WT control mice and M3Het animals increased as a function of CV*, where peak slow response amplitude was larger in irregular afferents as compared to regular afferents. However, in M3KO mice, the amplitude of EVS-mediated slow excitation decreased as a function of CV* (Fig 4B). The mean amplitude of EVS-mediated slow excitation in irregularly firing afferents (CV*>0.1) in M3KO mice (2.107±0.49, F(2,74)=21.12, p<0.0001, Dunn’s multiple comparison) was significantly different compared to controls (23.16±2.25) and M3Het animals (23.03±3.48, p<0.0001) (Fig 4C). In contrast, the mean amplitude of EVS-mediated slow excitation was similar in regularly-discharging afferents (CV*<0.1) from the three groups. (Control vs M3 Het vs M3 KO: 8.908±0.78 vs 6.91±0.99 (p=0.12) vs 7.04±0.84 (p=0.15), F(2,48)=0.2110, Tukey’s multiple comparison) (Fig 4D). EVS-mediated fast excitation (Fig. 4E), which is attributed to the activation of α4/β2 containing nAChRs (Holt et al., 2015; Schneider et al., 2021), was also similar between the three groups (Fast response amplitude>10sp/s: Control vs M3 Het vs M3 KO: 35.03±11.39 (n=7) vs 31.26±6.38 (n=14, p=.93) vs 26.6±6.24 (n=6, p=0.8), F(2,24)=0.4641, Tukey’s multiple comparison). The presence of EVS-mediated fast excitation confirmed that vestibular efferent neurons remain viable for electrical stimulation in M3KO animals.

### VsEP measurements in WT, M3 Het and M3 KO mice

Despite no obvious differences in background firing rates, M3KO mice had attenuated EVS-mediated slow excitation in irregularly-firing afferents but not regularly-firing afferents. We asked whether other functional response metrics might be altered by the loss of M3mAChRs. Vestibular sensory-evoked potentials (VsEPs) are compound action potential in response to linear acceleration of the head and they are thought to originate in irregularly-discharging calyx-bearing afferents (Curthoys et al., 2019; Curthoys et al., 2017; Jones et al., 2011; Jones et al., 2015; Lee et al., 2017; Ono et al., 2020). So, we recorded VsEPs in our control, M3Het, and M3KO animals to evaluate the functional impact of a blunted EVS-mediated slow excitation in irregular afferents. A typical VsEP recorded from a control animal is shown in Figure 5A where the P1N1 amplitude reflects the compound action potential from peripheral vestibular afferent neurons (Jones et al., 1999; Jones et al., 2004) while P2 and subsequent components reflect the activity of central vestibular relays within the brainstem and higher circuitry (Nazareth & Jones, 1998). Remarkably, VsEP metrics were indistinguishable across the three groups. P1N1 and P2N1 amplitudes (Fig. 5B), and P1, N1 and P2 latencies (Fig. 5C), as well as thresholds (Fig. 5D), were not significantly different among control, M3Het, and M3KO animals. These findings suggested that the irregularly-firing afferents in mice lacking M3mAChRs exhibit functional similarity to the control group, at least as far as basic VsEP metrics are concerned.

**Figure 5.**
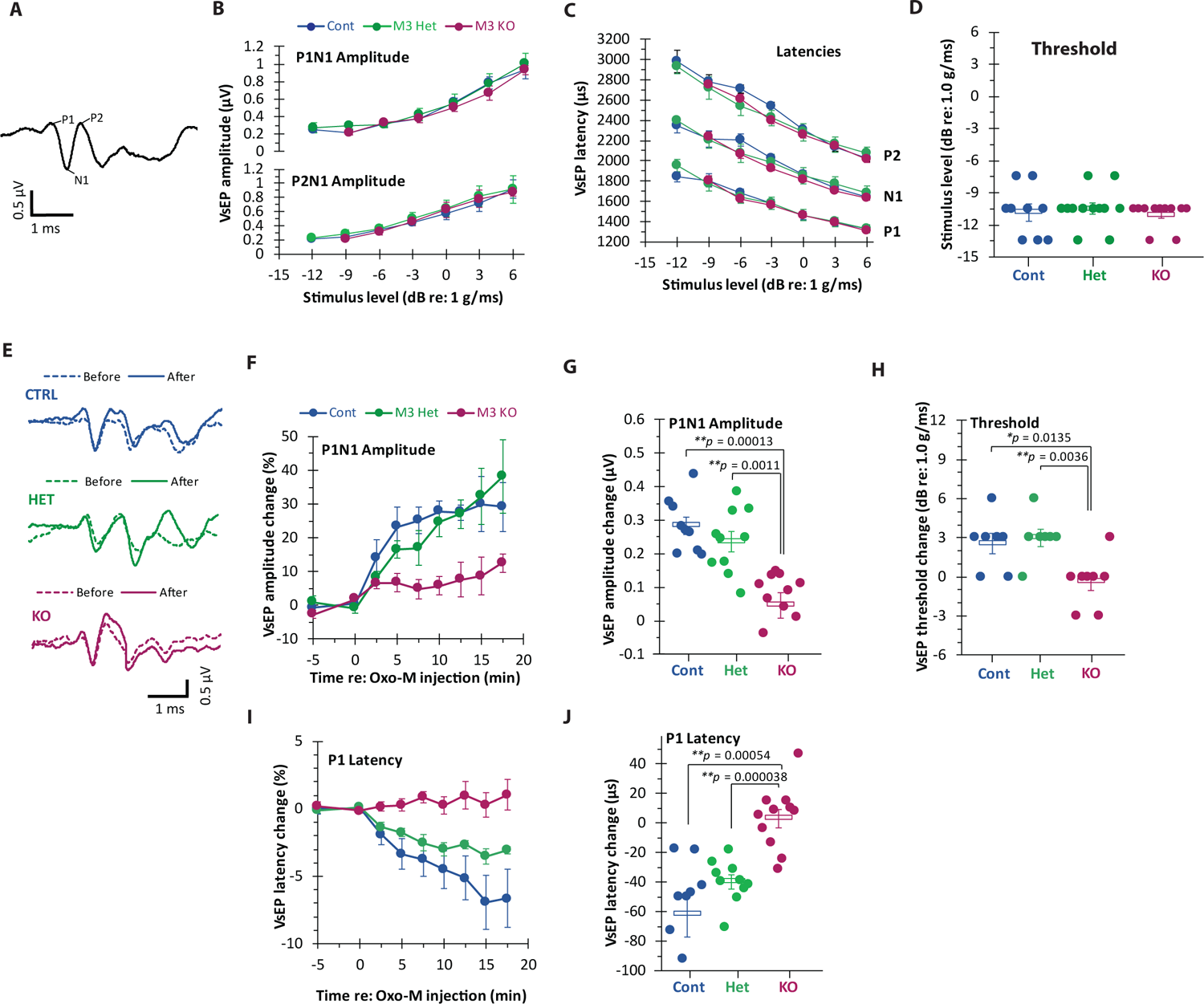
VsEP measurements in WT, M3mAChR-Het and M3mAChR-KO mice. **A**, Analagous to the ABR, the VsEP is a compound action potential generated in response to linear acceleration of the head. A mean VsEP trace is shown where positive and negative peaks are labeled. **B,** P1N1 and P2N1 amplitudes were not significantly different among wild type, M3mAChR-Het and M3mAChR-KO mice. **C,** P1, N1 and P2 latencies were not significantly different among wild type, M3Het, and M3KO mice. **D,** VsEP thresholds were not significantly different among wild type, M3Het, and M3KO mice. **E-J,** VsEP response metrics to the mAChR agonist, Oxotremorine-M (Oxo-M)**. E,** Representative mean VsEP traces before (dashed) and after IP administration of Oxo-M (solid). Significant enhancement in VsEP amplitude with time in WT and M3 mAChR Het mice but not in M3 mAChR KO mice. **F**, % change in P1N1 amplitude with IP-administration of Oxo-M in WT, M3 mAChR Het and M3 mAChR KO mice with time. Inset: Setup for VsEP measurement. **G,** Activation of mAChR by Oxo-M led to significant increase in VsEP amplitude of WT and M3 mAChR Het compared to M3 mAChR KO. **H**, Activation of mAChR leads to decrease in VsEP threshold of in WT and M3 mAChR Het compared to M3 mAChR KO. **I**, % change in P1 latency with time in WT, M3 mAChR Het and M3 mAChR KO mice. **J**, Activation of mAChR led to significant decrease in latency of P1 in WT and M3 mAChR Het compared to M3 mAChR KO.

We next investigated the effects of mAChR activation on VsEP metrics in the three groups of mice. Previous research has shown that afferent input sensitivity to mechanical or electrical stimuli is enhanced following the activation of afferent mAChRs by EVS stimulation or the application of the mAChR agonist Oxotremorine-M (Oxo-M) (Holt et al., 2017; Ramakrishna et al., 2021). We hypothesized that activation of mAChRs using, Oxo-M could be used to the effects of EVS stimulation on afferent mAChR activation. We predicted that application of Oxo-M would increase the P1N1 amplitudes of VsEPs as a reflection of the increase in peripheral afferent sensitivity. Consistent with this prediction, IP administration of 2mM Oxo-M in control mice enhanced P1N1 amplitudes over pre-drug values (Fig. 5E, 5F). P1N1 amplitude changes started as early as 2.5 mins following IP injection of Oxo-M and reached peak values around the 12.5-minute mark or so (Fig. 5F). Significant gradual P1N1 amplitude increases and P1 latency decreases were seen in the control (rmMANOVA, *p* < 0.001) and M3 Het (rmMANOVA, *p* < 0.0001) groups. The steady change in both P1N1 amplitude and P1 latency in the control and M3 Het group showed significant drug effects (post hoc Bonferroni, *p* < 0.01). At 7.5-15min post-Oxo-M administration, mean enhancement of P1N1 amplitudes by Oxo-M were significantly larger in controls (0.283 ± 0.027 µV; one-way ANOVA, *p* < 0.001) and M3Hets (0.237 ± 0.03µV; one-way ANOVA, *p* < 0.01), as compared with M3KO animals (0.047 ± 0.038 µV), consistent with the loss of M3mAChRs in irregular afferents from M3KO animals (Fig. 5G). Also, P1 latencies for controls (−62.176 ± 15.329 µs; one-way ANOVA, *p* < 0.01) and M3Hets (−39.91 ± 4.55 µs; one-way ANOVA, *p* < 0.01) were significantly shorter than those measured in M3KO animals (3.068 ±6.463μs) (Fig 5I, J).

We also monitored the effects of Oxo-M on response sensitivity to transient stimuli in control (N = 7), M3 Het (N = 7), and M3 KO (N = 8). In Fig. 5H, there was a small improvement in VsEP thresholds 10-15 mins following IP Oxo-M for both control (2.57 ± 0.78 dB) and M3 Het (3.00 ± 0.66 dB) groups, both which were statistically significantly different than the M3 KO group (−0.38 ± 0.68 dB). Hence, activation of mAChR on afferents can also potentially make afferents more sensitive to vestibular stimuli by lowering the afferent firing threshold. M3KO mice did not show this phenotype. Oxo-M administration did not significantly enhance VsEP P1N1 amplitude in M3 KO mice compared to the control group (Fig 5E-G). Additionally, M3 KO mice did not show a significant decrease in P1 latency or any significant changes in threshold which was seen in control mice (Fig 5H-J). These findings suggest that the changes seen in VsEP during Oxo-M administration is due to pharmacological activation of M3mAChRs on irregularly firing afferents, and mice lacking M3mAChRs did not have this response.

### Behavioral response of M3 mAChR KO mice to vestibular challenge

In lower vertebrates, EVS is thought to be involved in distinguishing between active and passive movement (Boyle & Highstein, 1990; Chagnaud et al., 2015; Straka & Chagnaud, 2017). Additionally, the EVS might be activated in response to somatosensory stimulation or to head-body rotation. In mammals, however, evidence suggests that EVS does not distinguish between active vs passive movement. Moreover, we don’t know specifically how EVS-mediated slow excitation affects vestibular behavior in mammals. To investigate this, we examined a panel of mouse behaviors in M3KO mice that can highlight changes in vestibular function including putative vestibulo-autonomic thermoregulatory responses, postural metrics, open field activity, gait analysis, balance beam test, and forced swim test. These assays are not exclusively vestibular but are impacted by peripheral vestibular alterations or deficits. We segregate these behavioral assays into those involving a vestibular challenge versus those simply measuring aspects of locomotion, navigation, and balance. We first wanted to evaluate the effects of vestibular challenge (VC) on thermoregulation and postural sway in M3 mAChR KO mice and their respective controls.

### Vestibular challenge induced thermoregulation

The mAChR nonselective antagonist, scopolamine is used to ameliorate the symptoms of motion sickness in human. Motion sickness, in human, leads to various phenotypic reactions including nausea, vomiting and hypothermia. Mice do not vomit but they experience motion-induced hypothermia and concomitant increases in tail temperature (Rahman & Luebke, 2023; Tu et al., 2017). This assay has been used to measure defects in presumed vestibular-autonomic function in mice lacking either the EVS neurotransmitter αCGRP or α9nAChRs, another postsynaptic target of the EVS. Mice lacking CGRP (Jones et al., 2018; Luebke et al., 2014) or the α9 nAChR (Chang et al., 2021; Hübner et al., 2015, 2017; Tu et al., 2017) show disruption of stereotypical thermoregulatory behavior. We wanted to test if M3mAChR KO mice also show a similar phenotype. For this assay, mice were rotated on an orbital shaker at 75rpm for 20mins while recording their body temperature (see methods). During the vestibular challenge (VC) control mice showed a characteristic drop in body temperature with a simultaneous increase in tail temperature (tail temp: Pre-VC vs during VC: 23.04±0.24 vs 27.62±1.91, n=6) which peaks at ∼9mins after the start of the rotation (Fig. 6A (top row), B). M3mAChR-KO mice did not show this rise in tail temperature (Fig 6A (bottom row), B) (Pre-VC vs during VC: 22.167±0.52 vs 22.55±0.39, n=3, p<0.001, rmMANOVA with post-hoc Bonferroni’s) although they still showed hypothermia due to rotation. The effect of both VC (F(25, 155)=2.816, p<0.0001) and genotype (F(1,7)=7.421, p=0.0296) were significant. Again, like the loss of CGRP and a9nAChRs, alterations in peripheral mAChR mechanisms appear to be associated with alterations in vestibular-mediated thermoregulatory changes.

**Figure 6.**
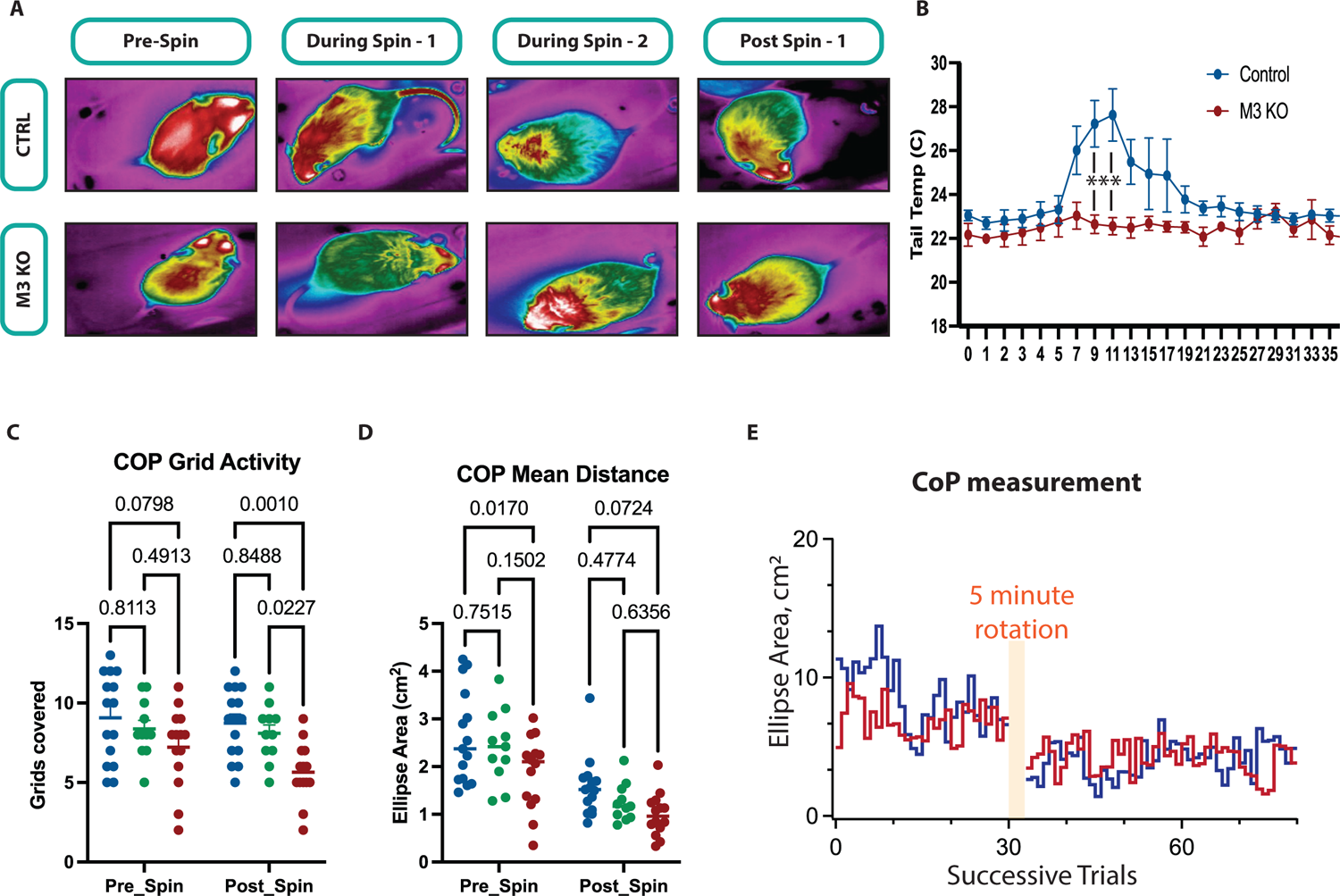
Behavioral response of M3 mAChR KO mice to vestibular challenge. **A, B**, Thermal changes to vestibular stimulation, Representative image of body temperature over the period of vestibular challenge of mice in the three groups. Pre-Spin: Thermal image before the start of vestibular stimulation; During Spin-1: Thermal image at 3min from the start of vestibular stimulation; During Spin-2: Thermal image at 10min from the start of vestibular stimulation; Post Spin: Thermal image at 10 min from the stop of vestibular stimulation. **B**, Graph representing the rise in tail temp with time for Wild type and M3mAChRKO. **C, D**, **Center of Pressure (Sway) changes in response to vestibular stimulation. C,** Graph showing the total number of grids covered before and after vestibular challenge in the wild type, M3mAChR Het and M3mAChR KO mice. D, Graph showing point to point distance covered by the animal before and after vestibular challenge in the wild type, M3mAChR Het and M3mAChR KO mice. E, Mean center of pressure (COP) measurements. The wild type (blue) and M3mAChR KO (red) mice do not have difference in CoP post vestibular challenge (VC).

### Changes in postural sway to vestibular challenge

Humans and mice exhibit small side-to-side or back-and-forth postural movements and adjustments to maintain our balance as we stand. These minute changes can be measured by having the subject stand on a force plate whose output identifies an elliptical sway space associated with postural adjustments. This sway space may vary depending on postural instabilities, vestibular dysfunction, or various neurological pathologies. We wanted to evaluate if M3mAChR-KO mice exhibit changes in postural sway following a vestibular challenge that differ from that seen in control animals. We calculated the sway path and center of pressure (CoP) for controls, M3Het, and M3KO mice as they stood on a small, enclosed force plate, before and after being rotated at 125 rpm for 5 mins on an orbital shaker. Ellipse area was calculated by using 95% confidence interval fit of the sway space. Mean distance travelled was calculated by calculating distance between the center of one ellipse to another. The number of grids covered before and after were not significantly different after VC, (Fig. 6C) (Grid activity: Pre-VS vs Post-VS: Control: 9.06±0.72 vs 8.73±0.55 (p=0.99); M3 mAChR Het 8.36±.054 (p=0.99), M3 MAChR KO 7.21±.0.67 vs 5.64±.0.48 (p=0.42), Tukey’s multiple comparison). The mean distance between the ellipse area center was used to estimate distance travelled by the mice. The mean distance pre- and post-VS was significantly different within the groups (Fig. 6D) (Pt to Pt distance: Pre-VS vs Post-VS: Control: 2.61±0.98 vs 1.57±0.62 (p=0.001); M3 mAChR Het 2.41±0.39 (p=0.99), M3 mAChR KO 1.87±.0.77 vs .98±.0.11 (p=0.42), Tukey’s multiple comparison). CoP area did not differ between control and M3mAChR KO mice post VC (Fig. 6E).

### Behavioral analysis of M3mAChR KO mice in absence of vestibular challenge

The limitation of our behavioral studies is that our mice model is a global constitutive M3mAChR KO (M3KO) animal. Hence some of the vestibular-related behaviors might not necessarily reflect loss of mAChRs in the peripheral vestibular system but instead could be attributed to mAChRs in other systems, particularly the CNS. However, it is important to characterize baseline behaviors in these animals and in instances of where there is convergence across multiple behavioral assays that might highlight changes in vestibular function. The effect of EVS-mediated slow excitation in awake, behaving mice has not been evaluated and our behavioral studies in the M3KO mice might reveal how the loss of mAChRs, and this EVS-mediated slow excitation, in the peripheral vestibular system impacts vestibular function. While M3KO mice showed normal VsEP amplitudes, threshold, and latencies suggesting normal-functioning irregular vestibular afferents innervating otolithic organs, we wanted to further evaluate other behavior including open field activity, gait, balance beam performance, and the forced swim test.

#### Open Field Activity

The vestibular system is involved in spatial navigation (Hufner et al., 2007; Laurens & Angelaki, 2020; Mao et al., 2021; Wallace et al., 2002; Yakusheva et al., 2007) and silencing or disrupting sensory information from vestibular periphery can lead to disruption in the formation of grid cells and head directions cells in mice (Angelaki & Laurens, 2020; Angelaki et al., 2020; Jacob et al., 2014; Laurens & Angelaki, 2018). Patients with bilateral vestibular damage also have atrophy in the hippocampus (Hufner et al., 2007). Additionally, transgenic mice with mechanotransduction defects from vestibular HCs also show deficits in open field activity (Asai et al., 2018; Pan et al., 2017; Wilkerson et al., 2018). To explore a possible role for EVS mediated slow excitation in spatial navigation, we performed open field activity on M3KO mice. M3KO mice has been shown to not have any movement deficit when evaluated over 48hrs using a photobeam interruptions (Yamada et al., 2001). For better tracking, we used high speed videography to track mice movements in a circular enclosure. Some key differences were noted among controls, M3Hets, and M3KO mice during the 10 mins within the enclosure. M3KO and M3Het mice covered less distance in the enclosure compared to control mice (Fig 7B) (Control (n=16) vs Het (n=10) vs KO (n=10): 38.31±2.13 vs 27.67±2.03 (p=0.0289) vs 22.25±2.3m (p=0.0002), two-way ANOVA with post-hoc Bonferroni), and exhibited lower mean speeds as well (Fig. 7C) (6.43±0.3 vs 4.63±.3 (p=0.0253) vs 3.7±0.3cm/s (p=0.0001), two-way ANOVA with post-hoc Bonferroni). Moreover, M3KO and M3Het mice were significantly less active compared to control (Fig. 7D) (Control vs M3 Het vs M3 KO: 68.47±2.7% vs 54.71±2.39% (p=0.0062) vs 54.2±3.57% (p=0.0037), two-way ANOVA with post-hoc Bonferroni). However, M3KO and M3Het were able to reach the comparable maximal speed which was not significantly different from control (Fig. 7E) (Control vs M3 Het vs M3 KO: 100.2±12.32 vs 122.1±27.92 (p=0.5913) vs 84.48±13.77 cm/s (p=0.7986), two-way ANOVA with post-hoc Bonferroni). To further isolate vestibular input for open field activity, we removed the contribution of visual input by performing the same experiment in the dark. We found that M3Het and M3KO mice were significantly less active in dark compared to control (Fig. 7D) (%time active: Control (n=16) vs M3 Het (n=24) vs M3 KO (n=16): 79.05±0.87 vs 61.39 ± 3.83 (p=0.0018) vs 45.13 ± 4.32 (p<0.0001), two-way ANOVA with post-hoc Bonferroni). Additionally, M3KO mice were also significantly less active compared to M3Het in dark (p=0.0022, two-way ANOVA with post-hoc Bonferroni). The presence of light (p=0.0096, F(1,33)=7.559, two-way ANOVA with post-hoc Bonferroni) and genotype (p<0.0001, F(2, 57)=15.55, two-way ANOVA with post-hoc Bonferroni) had strong effects on the active time spent in the open-field arena. M3KO and M3Het mice travelled significantly less distance in dark compared to control (Fig. 7B) (Control (n=16) vs M3 Het (n=24) vs M3 KO (n=16): 56.02±1.46 vs 43.42. ± 1.73 (p=0.0001) vs 30.1 ± 2.88 (p<0.0001), two-way ANOVA with post-hoc Bonferroni). Additionally, M3KO mice covered significantly less distance compared to M3Het in dark (p<0.0001, two-way ANOVA with post-hoc Bonferroni). The presence of light (p<0.0001, F(1,33)=56.94, two-way ANOVA with post-hoc Bonferroni) and genotype (p<0.0001, F(2, 57)=32.27, two-way ANOVA with post-hoc Bonferroni) had strong effect on distance travelled in the open-field arena. Similar to distance covered, M3KO and M3Het mice had significantly less speed in dark compared to control (Fig. 7C) (Control (n=16) vs M3 Het (n=24) vs M3 KO (n=16): 9.34±0.24 vs 7.25. ± 0.29 (p=0.0001) vs 5.02 ± 0.48 (p<0.0001), two-way ANOVA with post-hoc Bonferroni). Additionally, M3KO mice covered significantly less distance compared to M3Het in dark (p<0.0001, two-way ANOVA with post-hoc Bonferroni). The presence of light (p<0.0001, F(1,33)=55.16, two-way ANOVA with post-hoc Bonferroni) and genotype (p<0.0001, F(2, 57)=32.54, two-way ANOVA with post-hoc Bonferroni) had significant effect on speed of the mice in the open-field arena. Similar to the light condition, the three groups of mice did not significantly differ in their maximum speed in the dark (Fig. 7E) (Control vs Het vs KO: 51.96±6.13 vs 66.16±10.3 (p=0.5913) vs 41.71±5.73 (p=0.7986), two-way ANOVA with post-hoc Bonferroni). There was a significant effect of presence of light (p<0.0001, F(1,90)=22.88, two-way ANOVA with post-hoc Bonferroni) but not genotype (p=0.0573, F(2, 90)=2.953, two-way ANOVA with post-hoc Bonferroni) on maximum speed of the mice in arena. Thus, these set of experiment suggested that M3KO mice were further impaired in dark. Interestingly, the three groups didn’t significantly different in the % time spent in center vs periphery (data not shown) suggesting that anxiety phenotype might not be contributing to the phenotype we see here. Interestingly, we also observed some peculiar phenotypic difference in M3KO mice in open field activity where they seem to form activity hubs where they would return to during the course of experiment (Fig. 7E).

**Figure 7.**
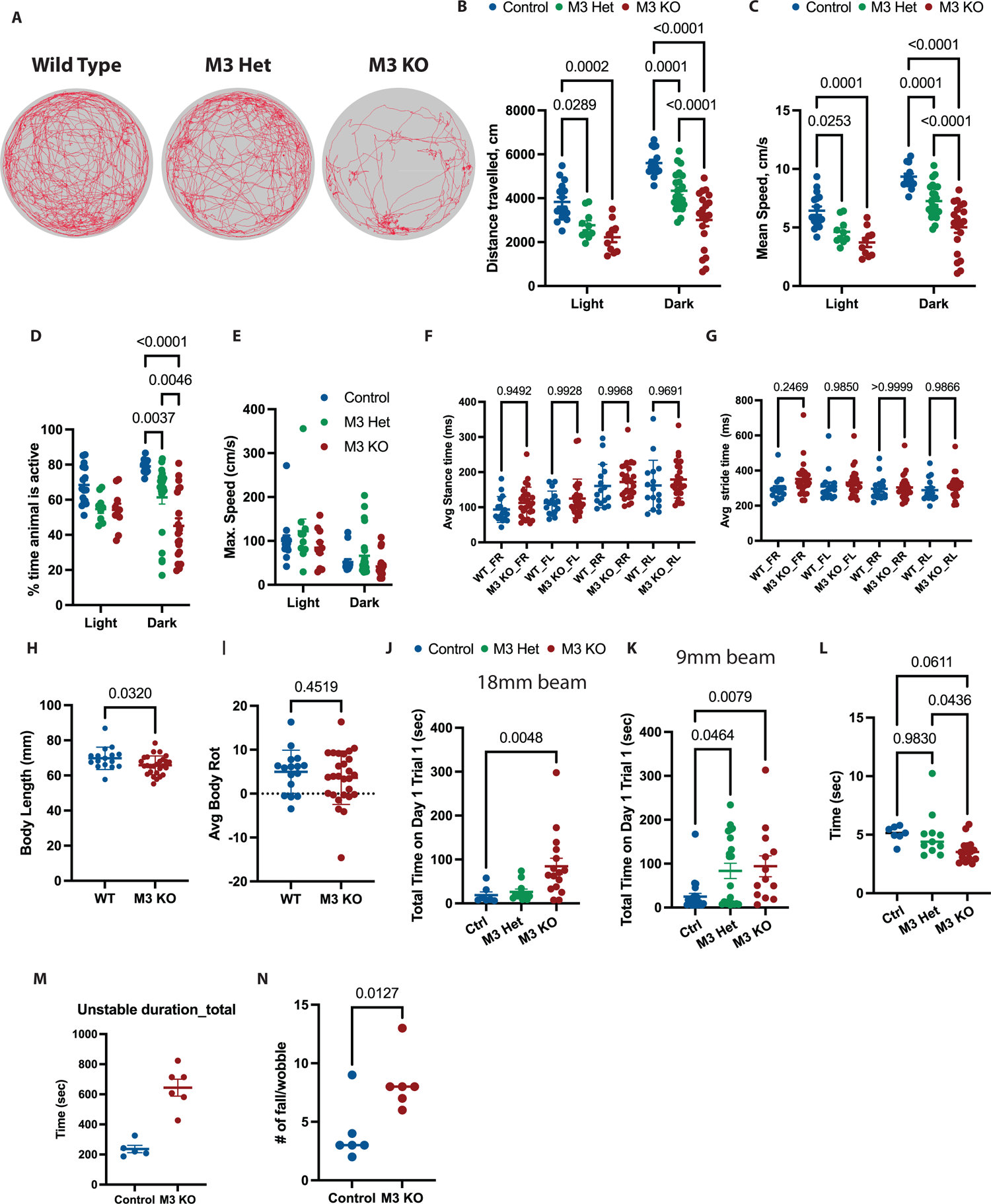
Behavioral analysis of M3 mAChR KO mice. **A-E,** *Open-field activity was measured in open circular arena (50” diameter) in light and in dark.* **A**, Example activity traces wild type, M3 mAChR Het (M3 Het) and M3 mAChR KO (M3KO) mice in light. **B,** Total distance travelled in 10mins in light and in dark by wild type, M3 mAChR Het and M3 mAChR KO mice. **C,** Mean velocity of wild type, M3 mAChR Het and M3 mAChR KO mice in light and in dark **D,** %time wild type, M3 mAChR Het and M3 mAChR KO mice were active in light and in dark. **E,** Maximum speed wild type, M3 mAChR Het and M3 mAChR KO mice could achieve in the open field arena in light and in dark. **F-I,** *Gait analysis,* Gait measurement were not significantly different between wild type and M3 mAChR KO mice. **J-L,** *Balance beam analysis,* **J-K**, Mice were trained either on 9mm or 18mm wide beam on Day 1 and Day 2. M3 mAChR KO mice took significantly longer to cross the beam on the training beam on Day1 Trial 1 compared to wild type and M3 mAChR Het mice when trained either 9mm (**J**) or 18mm (**K**). **L**, Mice trained on 20mm beam were tested on 9mm beam on Day 3. M3 mAChR KO showed no deficit in balance compared to wild type and M3 mAChR Het mice during the testing. **M-N,** *Forced swim test analysis.* **M**, Schematic of forced swim test timeline, mice underwent 6min (2 mins of training period and 4mins of testing period) of forced swimming and 10min of post-forced swim recovery. No significant difference was seen between wild type, M3 mAChR Het and M3 mAChR KO mice during the testing period. **N,** Post-forced swim test M3 mAChR KO mice were more unstable compared to WT and M3 mAChR Het mice as measured by duration of instability and number of falls. **O,** Post-forced swim test M3 mAChR KO mice also had more episodes of fall/wobble compared to WT and M3 mAChR Het mice.

#### Gait test

Open field activity analysis did not suggest any motor deficit as M3mAChR KO mice could reach the same maximum speed as the control mice (Fig 7C). Vestibular dysfunction is also commonly associated with gait disturbances and so we also evaluated the gait of control and M3KO animals. We looked at numerous parameters of gait in these mice and did not observe any significant difference in any metric between M3 mAChR KO mice compared to control (Fig 7F-I, one-way ANOVA with Tukey’s multiple experiment) (Avg: Average, Rot: Rotation, WT: Wild type, M3 KO: M3mAChR KO, FL: Front left, RR: Rear right, FR: Front right, RL: Rear left).

#### Balance Beam

Finally, we tested the balance of M3KO mice using a balance beam assay. For this, the mice were trained on either 18mm or 9mm wide beam for 3 trials/day for 2 days and finally tested them on day 3. M3mAChR KO mice took significantly longer time to cross the beam for the first trial on day 1 on both the 9mm and 18mm compared to control (Fig 7J-K) (Control vs M3Het vs M3KO: 9mm: 24.7±7.52sec vs 83.46±17.35sec (p=0.046) vs 94.08±23.88sec (p=0.0079): 18mm: 18.76±7.26 vs 26.17±6.45 (p=0.86) vs 88.44±18.3sec (p=0.0048), Kruskal-Wallis test with Dunn’s multiple comparison). However, the M3KO animals became proficient at crossing the beam and reaching the safe zone starting from trial 2 on Day 1. When mice trained on 18mm balance beam for 2 days and tested on 9mm on day 3, M3 KO mice took less time compared to WT control, but it was not significantly different (Fig. 7L) (Control vs M3 Het vs M3 KO: 5.09±0.25sec vs 4.97±0.6sec (p=0.9830) vs 3.65±0.24 sec (p=0.0611)).

Integrating the behavior of mice in open-field test with that on the balance beam, the M3KO mice seem to have issues making adjustments to a new environment or task but improve after some period of learning.

#### Forced Swim Test

To investigate the behavioral implication of proprioception in M3 mAChR KO mice. The mice were exposed to a forced swim test for 6mins followed by recovery under a heat lamp. The swim time was divided into 2min of training and 4min of testing. The M3KO mice performed no differently from the control WT mice or M3Het mice during the testing period (data not shown).

M3KO mice were not significantly different compared to control. However, we then observed something interesting in the M3KO mice during the recovery period. After transferring to the recovery area, M3KO mice had difficulty stabilizing their body and moving around the cage. The mice were repeatedly falling on their side or dragging their lower torso. We calculated the duration of instability in these mice as discussed in the method section. Compared to controls, M3KO mice had increased frequency of instability where they were either falling on their sides, dragging their body against the wall of the box or they wobbling as they were moving. The M3KO mice also took longer to recover from the instability, compared to controls, where a few M3 KO mice did not recover during the 10mins duration of observation (Fig 7M-N). This suggested that although the mice got accustomed to the swim environment during the testing period, they were not able to acclimate to the new environment when brought back to the recovery cage.

## Discussion

EVS mediated slow excitation is present in various species, including turtle, chinchilla, and squirrel monkey (see review (Jordan et al., 2015)) (Cullen & Wei, 2021; Holt, 2020). Moreover, it has been shown in several species that EVS-mediated slow excitation arises from activation of mAChR receptors (Holt et al., 2017; Schneider et al., 2021) though the specific receptor subtypes have not been explored. This has limited further exploration into understanding how EVS-mediated slow excitation is used to alter vestibular function. In this paper, we show that M3mAChRs are the main target of EVS-mediated slow excitation of vestibular afferents, particularly those with irregular discharge. Pharmacological data and residual EVS-mediated slow excitation in regular firing afferents suggest that M1mAChRs on the afferent may play a smaller role. It is currently unclear in the M3mAChR-KO animals whether the persistence of EVS-mediated slow excitation in regular afferents by M1mAChRs represents a simple unmasking or some compensatory mechanism engaged in the absence of M3mAChRs. Here, further testing in M1mAChR-KO animals might be instructive. We saw no evidence for M2 or m4mAChRs playing a role in EVS-mediated slow excitation either with selective pharmacological agents or in M2/M4-mAChR double KO animals. Finally, our pharmacological data with an M5mAChR PAM and NAM suggest that EVS-mediated activation of M5mAChRs can give rise to a much slower excitation. It is unclear where these putative M5mAChRs reside. Given its slower kinetics and no obvious interactions with the traditional EVS-mediated slow excitation, it may not be expressed in afferents but in hair cells instead where changes in transmitter release could enhance excitability of the afferent (Derbenev et al., 2005; Holt et al., 2003; Li & Correia, 2011; Yao et al., 2011).

In M3mAChR-KO mice, we expected that EVS-mediated slow excitation would be absent in vestibular afferents. However, regular firing afferents in M3KO mice still exhibited EVS-mediated slow excitation whose mean amplitude was not significantly different from control. However, we had not seen any difference in degree of pharmacological block between regular and irregular firing afferents by the M1mAChR antagonist telenzepine. The M1mAChR antagonist VU0255035 did exhibit a steady state blockade at the two lowest doses which might suggest blockade of M1mAChRs but the extent of the block did not seem to vary between regular and irregular afferents. A more detailed exploration is required. We think that the remaining EVS-mediated slow excitation in regular firing afferents of M3mAChR-KO(M3KO) mice might be a product of compensation. Recent experiments in the lab have been revisiting this issue by exploring the sensitivity of EVS-mediated slow excitation in M3KO mice to blockade by the M1mAChR antagonist, telenzepine. Our preliminary data suggest that EVS-mediated slow excitation in regular vestibular afferents of the M3KO mouse is more sensitive to telenzepine than regular afferents in control animals. In fact, in the few units we collected so far, a larger block of EVS-mediated slow excitation is seen at lower telenzepine concentrations, consistent with M1mAChRs taking over the process in compensating for lack of M3mAChRs. This compensation is common in the CNS where mice lacking M1 or M3mAChRs can be compensated by M3 or M1 mAChR, respectively. However, mice lacking both M1 and M3mAChRs are unable to compensate and have functional and behavioral defects(Chan et al., 2017). We would predict a complete loss of EVS-mediated slow excitation in M1/M3mAChR double KO animals.

Interestingly, mice lacking M3mAChRs had normal VsEPs suggesting that this receptor does not play a major role during afferent development at least as far as the mechanisms underlying discharge regularity are concerned. However, upon pharmacological activation of mAChRs using Oxo-M, evoked little or no enhancement of VsEP P1N1 amplitudes or a shift in P1N1 latencies or threshold in M3mAChR-KO mice consistent with loss of mAChRs on irregular afferents, as identified in our single-unit recordings from the same animals. The M3mAChR-KO mice provided us with a model to explore various vestibular behavior in animals where EVS-mediated slow excitation might be modulating irregularly-firing afferents.

One of the limitations of the behavior assays used in this paper is that M3KO mice we used are missing M3mAChRs in all cell types. These mice have been shown to have issues with digestion and smooth muscle contractility (Matsui et al., 2000; see review Matsui et al., 2004). We tried to counter these by supplementing IACUC approved special food with usual chow. M3KO mice also develop visual perception issues later in life (5+ month old: (Gericke et al., 2009; Gericke et al., 2011; Musayeva et al., 2021; Origlia et al., 2006)). We conducted our behavior experiments in mice aged between 2.5-4months to avoid any behavioral changes associated with visual deficits. M3mAChRs are also present in inhibitory neurons within the spinal cord that play a role as central pattern generator. However, we did not find any obvious deficits in gait in these mice. For future behavioral studies one could generate conditional M3mAChR KO mouse which lack M3 mAChR in vestibular afferents only. This would eliminate mAChR issues elsewhere and focus on behavioral consequences of EVS-mediated slow response in isolation. However, currently there is a lack of genes unique to vestibular afferent that we could currently leverage. Although the lack of M3mAChRs in the CNS that might be affecting our observations, we were able to look at various vestibular behaviors in these mice to see if any obvious functional changes could be observed.

We show that mice lacking M3mAChRs seem to behave differently in a number of behavioral assays. We noted that these mice behaved quite differently when first introduced to new environments (open field activity arena), elevation (balance beam), water (swim test), and return to warming arena after swim test, though they neither exhibited movement deficits or obvious balance issues. They also had issues in vestibular-mediated thermal shifts, similar to what is seen in mice lacking other transmitters or receptors involved in EVS-afferent interactions (Chang et al., 2021; Hübner et al., 2015, 2017; Jones et al., 2018; Luebke et al., 2014; Tu et al., 2017). These mice clearly had severe balance issue after the forced swim test which could be seen in the warming recovery arena where they drag their tails, fall on their sides, and/or cling to the cage wall to stabilize/move around. These observations could not be explained by movement deficits alone.

## Acknowledgment

This research was supported by NIH/NIDCD Grants R01DC0016974 (JH). We thank Dr. Anne Luebke at University of Rochester for providing equipment to use for behavioral experiments. We would also like to thank Dr. Shafaqat M Rahman, Abigail Dweh and Carissa Minarovich for their help with thermal experiments. We thank URMC Behavioral Sciences Facility Core for their help with Gait and Forced Swim test experiments. We would also like to thank Dr. Dr. Jürgen Wess at NIH/NIDDK for providing us with M2/M4 mAChR DKO and M3 mAChR KO mice.

---

**Supplementary Figure 7.**
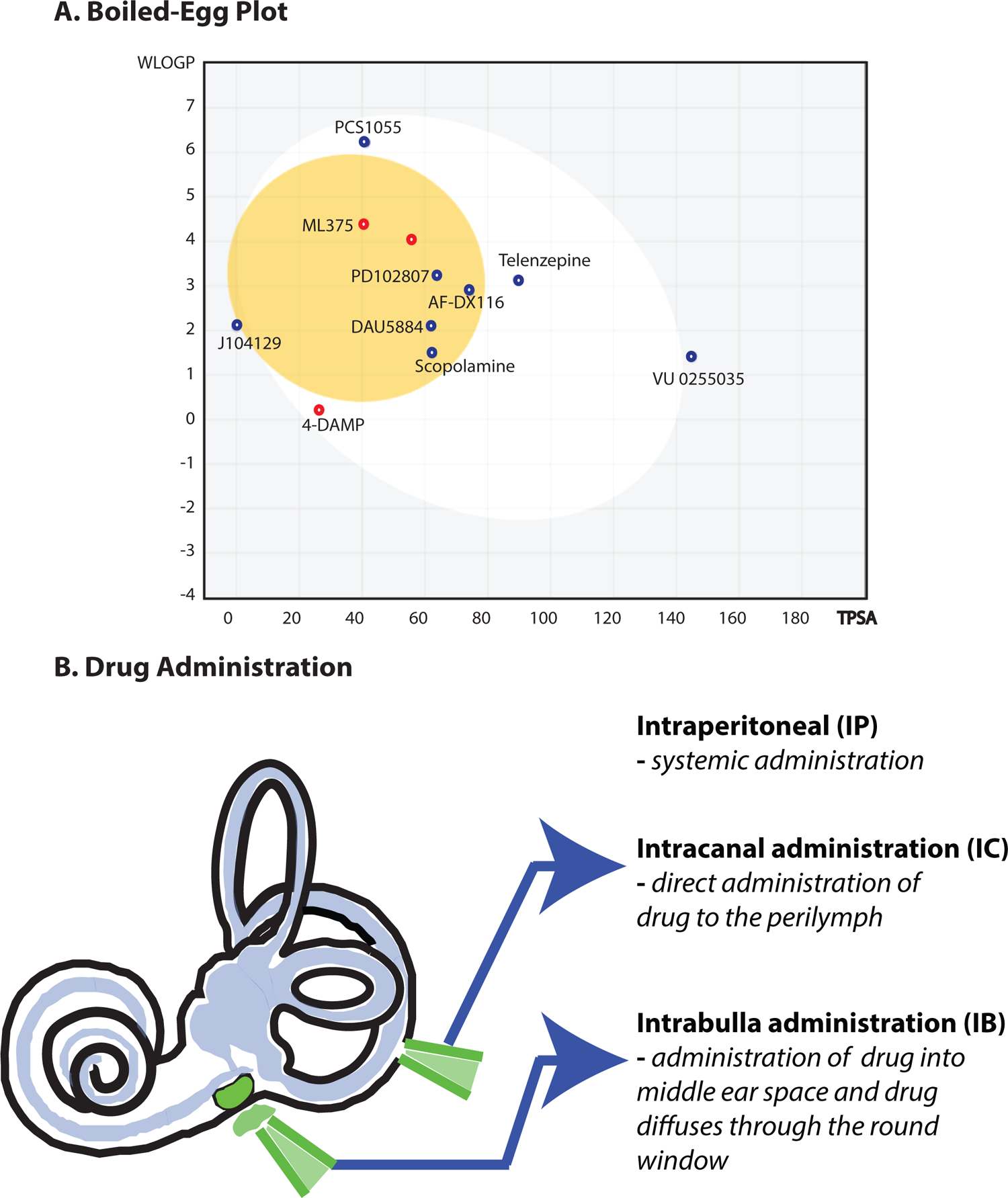

